# Ubiquitin selective ribosome profiling reveals systematic principles of co-translational quality control

**DOI:** 10.64898/2026.06.18.733060

**Authors:** Maximilian Seidel, Shengdi Li, Katharina Geissler, Xavier Hernandez Alias, Iskander Khusainov, Zinaida Kucherenko, Anja Becker, Daphne K. Welther, Bahtiyar Kurtulmus, Jan Provaznik, Jie Wu, Julian D. Langer, Danny Nedialkova, Julien Gangeur, Vladimir Benes, Mikhail M. Savitski, Judith Frydman, Martin Beck, Lars M. Steinmetz

## Abstract

Protein biogenesis is a stress- and error-sensitive process that can lead to nascent protein misfolding and aggregation, challenging cellular proteostasis.^1^ Co-translational ubiquitination (CTU) is a critical surveillance mechanism,^2,3^ yet its regulatory principles remain unclear due to limited known substrates and the lack of translatome-wide methods to query CTU. Here, we introduce UbSeRP, an approach that enables ubiquitin linkage-specific, and translatome-wide mapping of CTU. Focusing on ribosome-associated quality control (RQC), we expand the known endogenous RQC substrates in *S. cerevisiae* from few^4–7^ to thousands, reveal that only a subset of all identified disomes undergoes RQC, and uncover biophysical features of nascent chains that predict RQC engagement. We also identify widespread RQC-independent ubiquitination of nascent proteins, implicating broader roles of CTU in protein complex assembly. In a chronological aging model, we show that aging remodels translation and diminishes RQC engagement, favoring RQC-independent proteasomal pathways. Our findings provide systematic insight into the determinants and adaptability of CTU in maintaining proteostasis under changing physiological conditions.

## Main

Correct protein biogenesis is essential to maintain a functional proteome.^8^ Consequently, errors during translation that result in aberrant nascent chains contribute to proteotoxic stress and proteostatic decline. Cells therefore rely on quality control mechanisms that detect and resolve aberrant translation states, ideally while the polypeptide is being synthesized.^9^

Ubiquitination has emerged as a central regulator of co-translational quality control.^2,3,10,11^ Ribosome stalling and collisions, resulting in the formation of disomes, can trigger co-translational ubiquitination (CTU) of both the ribosomal proteins and the associated nascent chains by the ribosome-associated quality control (RQC) machinery (CTU_RQC_). This leads to ribosome disassembly and nascent chain degradation.^12^ However, CTU can also occur on nascent chains (NC) independent of RQC (CTU_NC_), for example mediated by the human E3 ubiquitin ligase TRIM25,^13^ indicating broader roles for CTU beyond RQC surveillance, such as clearing of defective folding or assembly intermediates.^2,3,10,11,13^

Progress in understanding CTU has been limited by technical constraints. Only a small number of endogenous RQC substrates have been identified.^4–7^ More importantly, previous studies did not distinguish CTU_RQC_ from CTU_NC_ in a translatome-wide manner^3,14,15^ and therefore, the prevalence of the two CTU modes remains unclear. Moreover, it is not understood how these CTU pathways adapt to physiological challenges such as aging, a condition proposed to result in a RQC overload.^16,17^

Here, we develop ubiquitin selective ribosome profiling (UbSeRP), an approach enabling translatome-wide, site-specific detection of CTU under endogenous conditions. Applying UbSeRP in *Saccharomyces cerevisiae*, we systematically map CTU_RQC_ substrates and uncover distinct characteristics that link nascent chain biophysical properties and disome spacing to CTU_RQC_ and uncover that only a fraction of all disomes engage in CTU_RQC_. We additionally identify widespread CTU_NC_ within and proximal to protein domains associated with protein complex assembly, revealing an unexpected role for CTU during translation. Finally, by integrating UbSeRP with proteomics of ribosome-nascent chain complexes in aged yeast cells, we show that aging alters translation elongation and shifts the balance of CTU pathways. Together, our results establish CTU as a pervasive and adaptable regulator of co-translation quality control across cellular states.

### An experimental framework to characterize CTU

UbSeRP is based on the combination of three analyses (**Fig. 1a**). First, we analyzed monosome (26–34 nt) and disome (54–68 nt) footprints subsequent to ubiquitin-affinity enrichment of ribosome-nascent chain complexes (RNCs). Second, we used linkage-selective TUBEs,^18^ including a PAN-ubiquitin (PAN-Ub) probe that recognizes polyubiquitin chains regardless of linkage type, as well as K48- or K63-ubiquitination specific probes. Mock enrichments using empty beads incubated with RNCs were used to correct for non-specific binding by subtracting mock-enriched ribosome profiles from TUBE-enriched samples. Third, we compared cycloheximide (CHX) and puromycin-treated lysates to distinguish CTU_NC_ from CTU_RQC_. Under high-salt lysis conditions, puromycin partially releases nascent chains without disrupting ribosomes,^19^ whereas cycloheximide stabilizes RNCs. While monosome versus disome analysis provides a means to distinguish translating from stalled ribosomes, reads that are diminished in puromycin compared to cycloheximide experiments indicate ubiquitination of the nascent chain instead of the ribosome. To confirm robust enrichment and data quality, we performed orthogonal biochemical and sequencing analyses, which validated the specificity and reproducibility of UbSeRP (**Supplementary Figs. 1, 2**).

**Figure 1.**
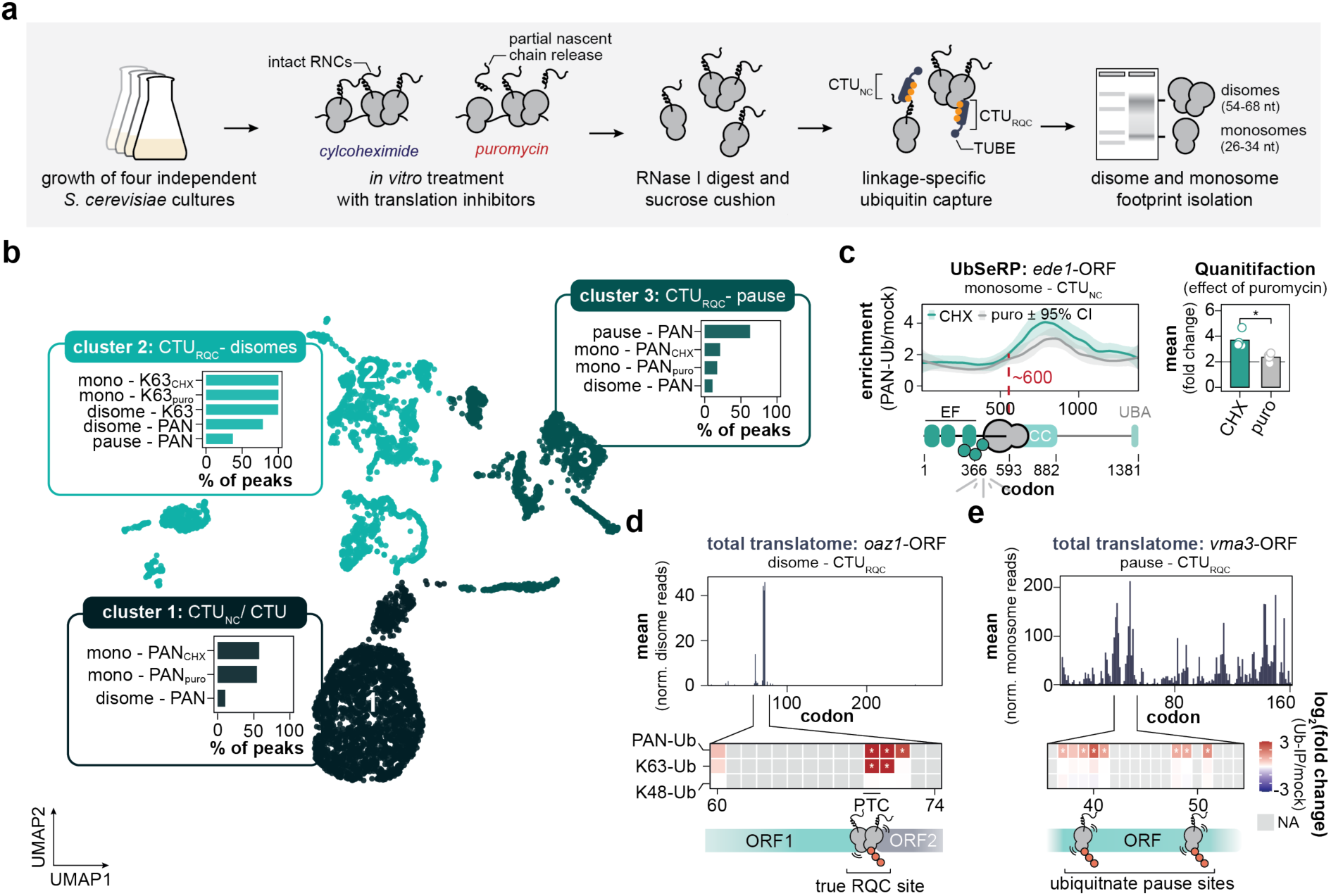
Ubiquitin selective profiling recapitulates true-positive targets of CTU. **a,** Schematic overview of the ubiquitin selective ribosome profiling (UbSeRP) workflow in *S. cerevisiae*. **b,** UMAP of significant UbSeRP peaks across the different UbSeRP modalities. Bar graphs display most prominent datasets per cluster (each having more than 100 significant peaks). **c,** Selective ribosome profile for *ede1*-ORF obtained from PAN-Ub pulldowns (*left*) and a bar plot of the enrichment quantification using the mean log_2_(fold change) (*right*). Nascent Ede1 is enriched at codon ∼600. Profiles depict LOESS-smoothed enrichment profiles (PAN-Ub enrichment *versus* mock non-binding control) for CHX- and puro-treated ribosome profiling experiments from four biologically independent experiments (*n*=4). The shaded area indicates the bootstrapped 95 % confidence interval (CI). \**p* = 0.011 (cycloheximide *versus* puromycin for PAN-Ub enrichment, two-sided paired t-test). **d,** Bar plot of mean normalized total disome reads for *oaz1*-ORF. Below, heatmap of Ub-enrichments (DESeq2-analysis). log_2_(fold change) scale is limited to 3 and -3; codon positions with a significant adjusted *p*-values are displayed as asterisk (\**padj* ≤ 0.05). NA (not applicable) corresponds to codon sites where log_2_(fold change) was not determined due to low reproducibility or expression cutoffs. Graph displays UbSeRP data from four biologically independent experiments for each condition (*n*=4). **e,** same as in **d,** but for *vma3*-ORF as a representative example of CTU_RQC_-pause. Note, *vma3*-ORF is exclusively targeted by CTU_RQC_-pause. CHX: cycloheximide, ORF: open reading frame; PTC: premature stop codon. puro: puromycin; TUBE: tandem ubiquitin binding entity; Ub: ubiquitin; FC: fold change. n.s., *p* > 0.05; \**p* ≤ 0.05; \*\**p* ≤ 0.01; \*\*\**p* ≤ 0.005; \*\*\*\**p* ≤ 0.001.

To resolve distinct CTU pathways, we developed a statistical framework to analyze our multimodal UbSeRP data to derive CTU_NC_ and CTU_RQC_ for pause and disome sites, respectively (**Supplementary Fig. 3a**). Using the significant ubiquitin-enriched regions within the translatome, we generated a uniform manifold approximation and projection (UMAP), which revealed three main clusters (**Fig. 1b**). Cluster one was enriched for monosome peaks with puromycin-sensitivity (**Supplementary Fig. 3b**), consistent with CTU_NC_. This includes PAN-Ub peaks of the *ubi4*-ORF that encodes a polyubiquitin precursor^20^ and serves as an internal reference for ubiquitin enrichment and puromycin-sensitivity rather than a ubiquitinated substrate (**Supplementary Fig. 3c**). Guided by these puromycin-sensitive signatures, we extended our framework with a dedicated pipeline inspired by Bertolini et al.^21^ to detect CTU_NC_ in monosome profiling data based on differential enrichment between CHX and puromycin conditions (**Supplementary Fig. 3d-f; Methods**). This revealed nascent chain ubiquitination onset sites, for example within Ede1 (**Fig. 1c**).

Cluster two was defined by disome and pause peaks. It contained the highest number of K63-Ub positive peaks across different categories, consistent with this cluster representing CTU_RQC_ targets.^22,23^ Within this cluster, we retrieved the two known RQC substrates *oaz1* (**Fig. 1d**) and *sdd1* **(Supplementary Fig. 3g)**, with sharp disome peaks, as well as PAN- and K63-Ub enrichment at known RQC motifs.^4,5^

In cluster three, we found translation elongation pause sites (**Fig. 1e**), where stalled ribosomes have not formed detectable disomes, suggesting that these paused ribosomes may engage in different quality control pathways. Together, UbSeRP can distinguish endogenous CTU pathways and provide a framework for the downstream analyses. In the following, we will define the mechanistic and biological signatures of CTU_RQC_ (**Figs. 2,3**) and CTU_NC_ (**Fig. 4**) and their context dependence in chronological aging (**Fig. 5**).

**Figure 2.**
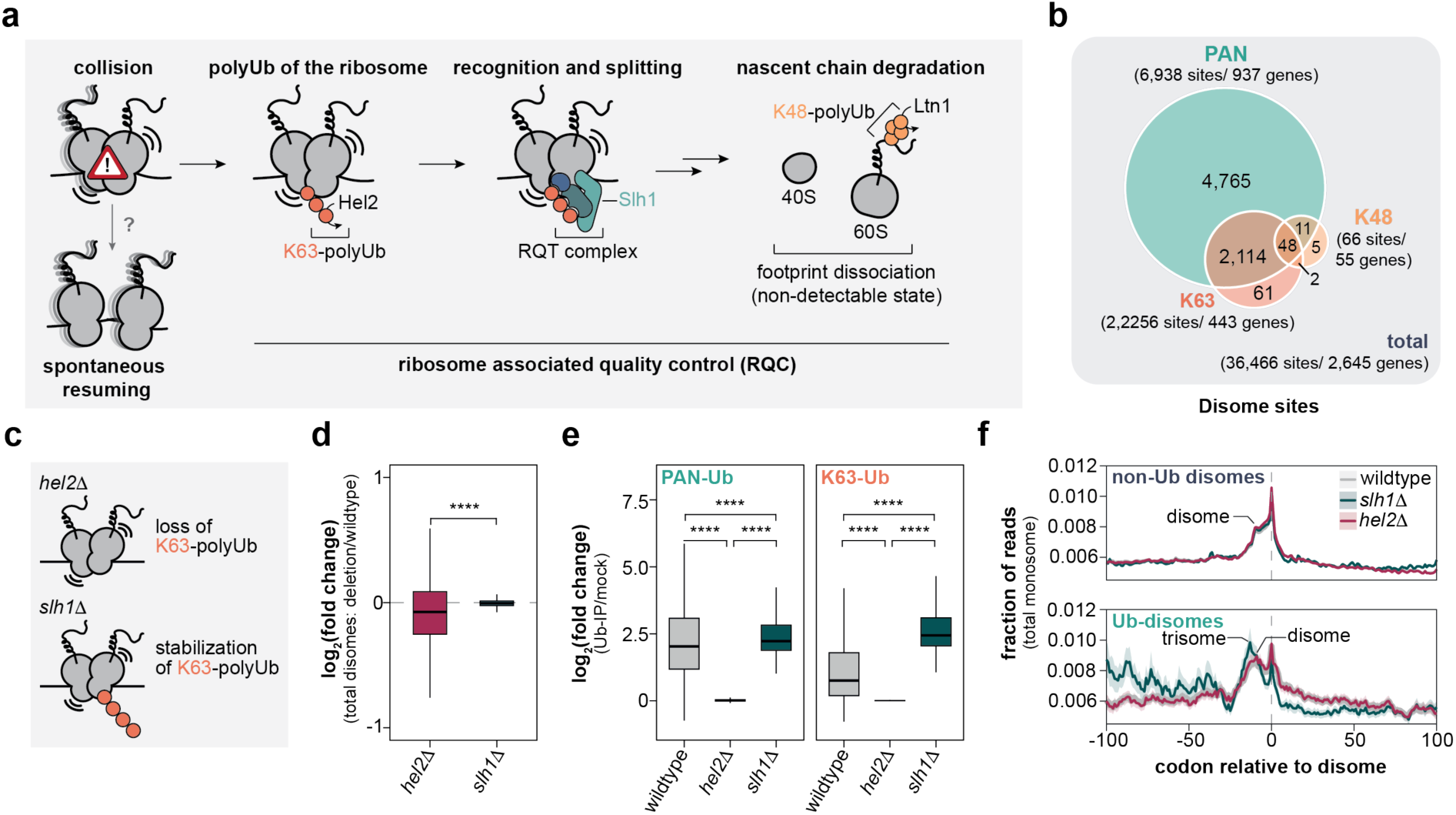
UbSeRP reveals RQC targets. **a,** Schematic representation of the ribosome-associated quality control (RQC) pathway adapted from Inada.^12^ **b**, Venn diagram showing the overlap of disome sites according to their ubiquitination status. Total disomes represent sites reproducibly identified in 3 of 4 replicates with more than 10 normalized reads. Ubiquitinated disomes were defined as sites with a log_2_(fold change) ≥ 1 and an adjusted *p*-value (*padj*) ≤ 0.05. **c,** Schematic of the expected effect of *hel2Δ* and *slh1Δ* on the ubiquitination of disomes. **d,** Box plot of the log_2_(fold change) of total disomes comparing RQC-mutants to wildtype (*n*= 36,466). **e,** Box plots showing the log_2_(fold changes) for ubiquitinated disome sites found in *slh1*Δ (log_2_(fold change) ≥ 1, *padj* ≤ 0.05, PAN, *n*=2,110; K63, *n*=2,222) in comparison to wildtype and *hel2*Δ experiments for the same sites. For **d** and **e**, the central line in the box plot depicts the median, box edges represent the interquartile range (IQR), and the whiskers represent the 1.5 IQR. \*\*\*\**p*= 0 (PAN- or K63-Ub for wildtype or *slh1*Δ *versus hel2*Δ), \*\*\*\**p*= 3.318×10^-15^ (K63-Ub for wildtype *versus slh1*Δ), \*\*\*\**p*= 1.295×10^-285^ (K63-Ub for wildtype *versus slh1*Δ, Wilcoxon test). **f,** Metagene plot of monosomes at non-ubiquitinated (*top*) and ubiquitinated disome sites (*bottom*). (non-ubiquitinated disome sites *n*= 30,118, ubiquitinated disome sites, *n*= 2,094). Solid line represents the mean fraction of reads and shaded area the 95% CI. n.s., *p* > 0.05; \**p* ≤ 0.05; \*\**p* ≤ 0.01; \*\*\**p* ≤ 0.005; \*\*\*\**p* ≤ 0.001.

**Figure 3.**
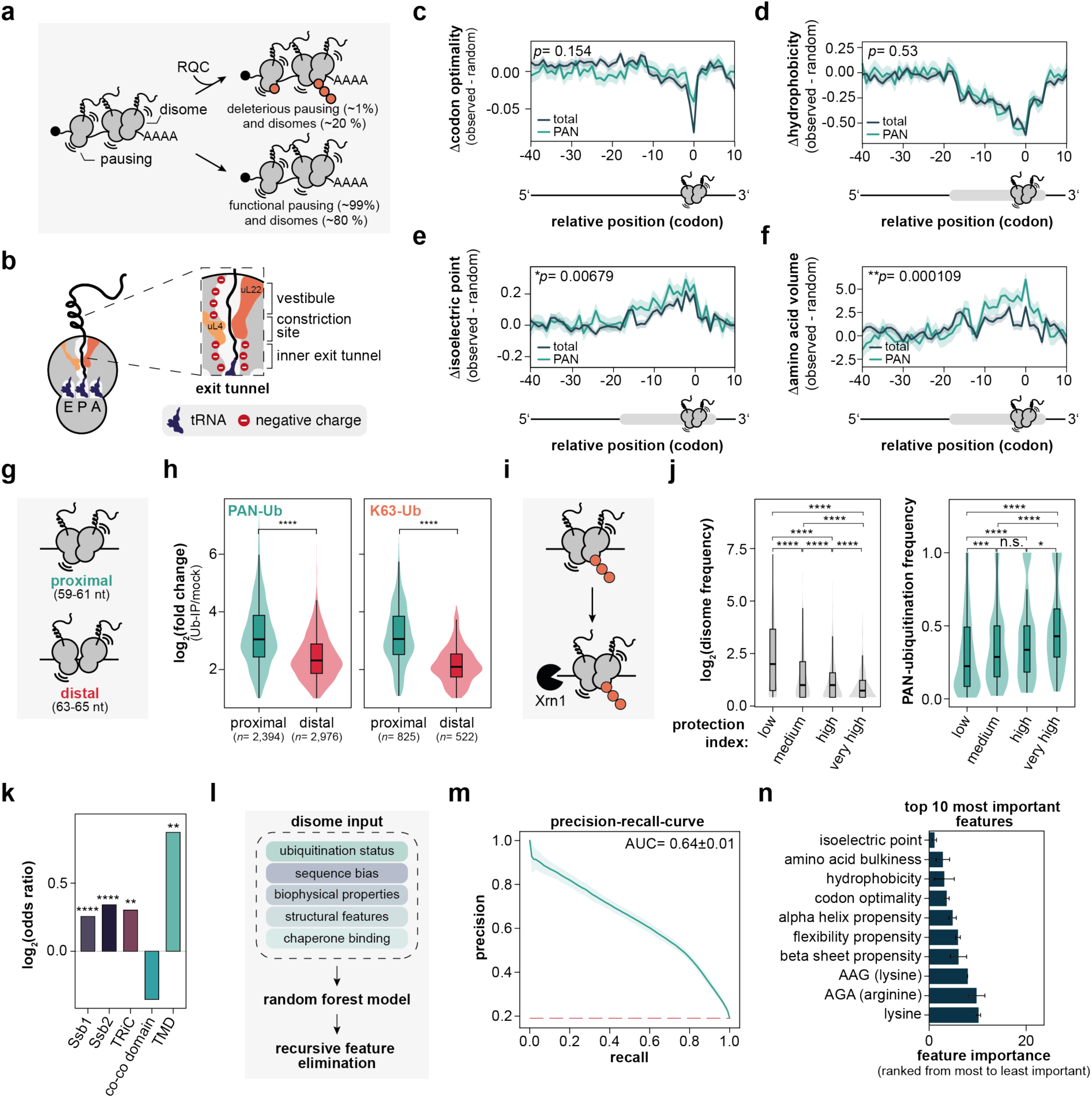
UbSeRP reveals mechanistic insights into substrate-selectivity of RQC. **a,** Schematic representation of translational pausing and disome formation leading to RQC, distinguishing functional pausing from deleterious disomes. **b,** Schematic of the ribosomal exit tunnel. The negatively charged ribosomal exit tunnel narrows approximately 30 Å upstream of the P site. Figure adapted from ^36^. **c,** Metagene plot of codon optimality at disome sites. The solid line indicates the mean Δcodon optimality (observed minus background) at disome sites, and the shaded area denotes the 95% confidence interval. No significant difference was observed between PAN-ubiquitinated disomes and total disomes (Wilcoxon test, *p*= 0.154). **d–f,** Metagene plots showing changes in hydrophobicity (**d**), isoelectric point (pI; **e**), and amino acid volume as a measure of bulkiness (**f**) at disome sites. In each panel, the solid line indicates the mean change at disome sites and the shaded area denotes the 95% confidence interval. Statistical significance was assessed using the Wilcoxon test: hydrophobicity, *p*= 0.53 (n.s.); pI, \*\**p*= 0.00679; bulkiness, \*\*\*\**p*= 0.000109 for PAN-ubiquitinated disomes versus total disomes. **g,** Schematic representation of proximal and distal disomes and their corresponding footprint bins. **h,** Violin plots of log_2_(fold changes) for Ub-IP/mock for PAN-Ub (*left*) and K63-Ub (*right*). Center lines indicate the median, box boundaries indicate the interquartile range (IQR), and whiskers extend to 1.5 IQR. Statistical significance was assessed using the Wilcoxon test: PAN-Ub, \*\*\*\**p*= 8.21×10^-221^ for proximal *versus* distal disomes; K63-Ub, \*\*\*\**p* = 8.21×10^-221^ for proximal *versus* distal disomes. **i,** Model illustrating the intersection of RQC and co-translational mRNA decay mediated by the 5′–3′ exonuclease Xrn1. **j,** Correlation of disome frequency and PAN-ubiquitination frequency with the codon protection index reported by Pelechano et al.^34^ Center lines indicate the median, box boundaries indicate the IQR, and whiskers extend to 1.5 IQR. Low codon protection index corresponds to low mRNA degradation, very high protection strong levels of degradation. For disome frequency, group sizes were: low, n = 757; medium, *n=*648; high, *n=*539; very high, *n=*473. For PAN-ubiquitination frequency, group sizes were: low, *n=*331; medium, *n=*216; high, *n=*296; very high, *n=*40. Pairwise comparisons of collision frequency for different codon protection bins were performed using the Wilcoxon test: low *versus* medium, \*\*\*\**p*= 1.22×10⁻²⁰; low *versus* high, \*\*\*\**p*= 1.12×10⁻³²; low *versus* very high, \*\*\*\**p*= 2.55×10⁻⁴⁸; medium *versus* high, \*\*\*\**p*= 4.66×10⁻⁴; medium *versus* very high, \*\*\*\**p*= 9.97×10⁻¹³; and high *versus* very high, \*\**p*= 3.03×10⁻⁴. PAN-ubiquitination frequency across codon protection bins was compared pairwise using the Wilcoxon test: low *versus* medium, \*\*\**p*= 2.79×10⁻³; low *versus* high, \*\*\*\**p*= 8.91×10⁻⁶; low *versus* very high, \*\*\*\**p*= 4.08×10⁻¹¹; medium *versus* high, n.s., *p*= 7.63×10⁻²; medium *versus* very high, \*\*\**p*= 3.25×10⁻⁵; and high *versus* very high, *p*= 3.41×10⁻². **k,** Enrichment analysis of chaperone binding and structural features. Chaperone binding and transmembrane domains (TMDs) are enriched among RQC-associated disomes, whereas co-co-translational assembly domains (co-co domains) are associated with non-ubiquitinated disomes. Chaperone binding sites and structural features were obtained as detailed in **Methods**. Statistical significance was assessed using Fisher’s exact test: Ssb1, \*\*\*\**p* = 1.06×10^-9^; Ssb2, \*\*\*\**p* = 1.16×10^-14^; TRiC, \*\**p*= 1.08×10^-3^; co-co-translational assembly domain, n.s., *p*= 0.107; TMD, \*\**p*= 1.03×10^-3^. **l,** Schematic of the random forest classifier. **m,** Precision–recall curve showing recovery of RQC events among all identified disomes. The shaded area indicates ±1 standard deviation across 100 cross-validation folds (10 repeats for 10-fold stratified CV). The red dashed line indicates the expected precision of a random classifier (class prevalence baseline). **n,** Bar plot showing the ranking of features by Recursive Feature Elimination (RFE), ordered from most to least important (rank 1 = most important, *top*). Error bars indicate standard deviation of the mean RFE rank across 10 stratified cross-validation folds. n.s., *p* > 0.05; \**p* ≤ 0.05; \*\**p* ≤ 0.01; \*\*\**p* ≤ 0.005; \*\*\*\**p* ≤ 0.001.

**Figure 4.**
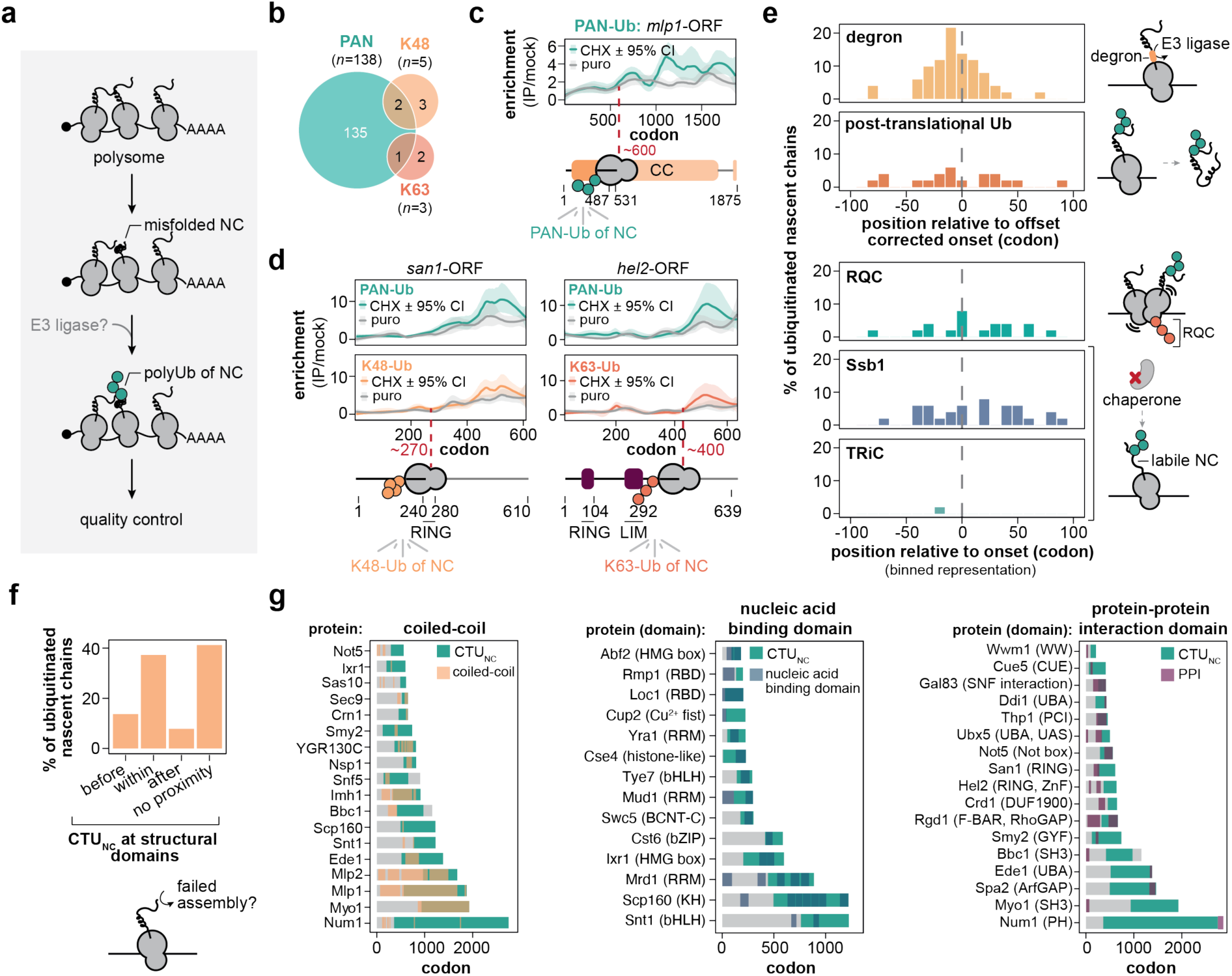
UbSeRP reveals RQC-independent ubiquitination of nascent chains. **a,** Schematic illustrating the impact of co-translational ubiquitination of nascent chains (CTU_NC_) on potential quality control. **b,** Venn diagram showing the identified nascent protein targets prior to manual curation. **c,** Representative ribosome profiles for the PAN-Ub target *mlp1*-ORF. Plots show LOESS-smoothed UbSeRP enrichment (PAN-Ub *versus* mock) from cycloheximide (CHX)- or puromycin (puro)-treated ribosome profiling experiments across four biologically independent replicates per condition (*n*=4). Shaded areas indicate bootstrapped 95% confidence intervals. Red dashed lines mark the inferred ubiquitination onset, defined by an enrichment threshold of 2. **d,** same as in **c**, but for *san1-*(PAN- and K48-Ub-enrichments *versus* mock; *left*) and *hel2-*ORF (PAN- and K63-Ub-enrichments *versus* mock; *right*). **e,** Histogram showing the distribution of degrons, ubiquitinated lysins, RQC sites, Ssb1-binding sites, and TRiC-binding sites relative to nascent chain ubiquitination onsets (*left*), with a schematic summary (*right*). For degrons and ubiquitinated lysins, a 30-amino-acid offset was applied to account for delayed exposure caused by the ribosomal exit tunnel. No such correction was applied to RQC, Ssb1, or TRiC sites, as these positions were derived from SeRP experiments. Degron annotations were obtained from Degronopedia,^39^ ubiquitinated lysins from Swaney *et al.*,^38^ RQC sites from this study; Ssb1 sites from Döring *et al*.,^40^ and TRiC sites from Stein *et al*.^41^ **f,** Bar plot showing the proportion of ubiquitination onsets located within a structural domain, within a 30-amino-acid window before or after a domain boundary and no domain association. **g,** Heatmap showing the overlap between coiled-coil domains (*left*), nucleic acid-binding domains (*middle*), and protein–protein interaction domains (*right*) with co-translational ubiquitination of nascent chains (CTU_NC_). Only proteins included in the manually curated target list were analyzed. CHX: cycloheximide, CTU_NC_: CTU of nascent chain; ORF: open reading frame; puro: puromycin; Ub: ubiquitin.

**Figure 5.**
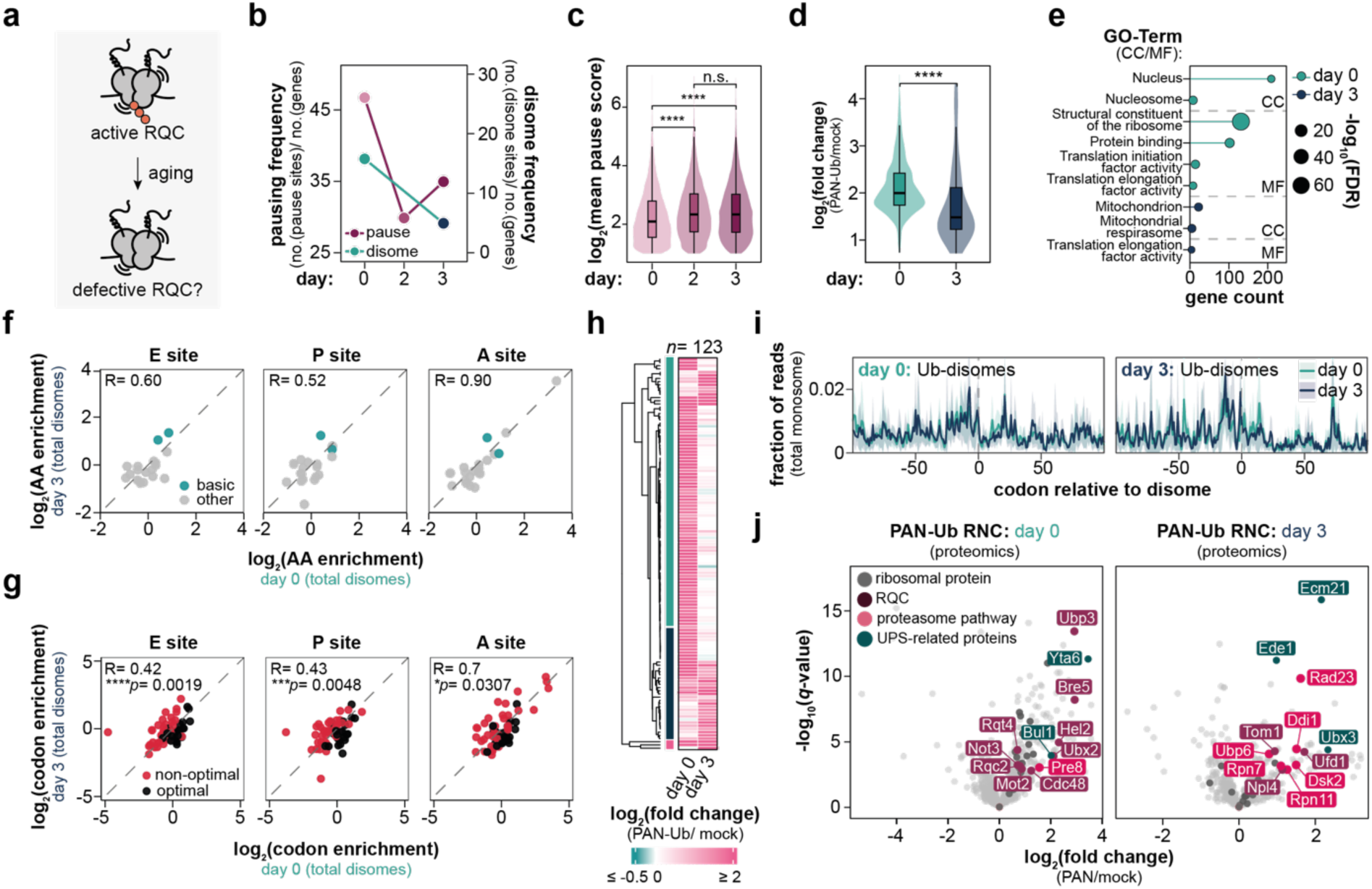
Age-associated remodeling of CTU during chronological aging in *S. cerevisiae*. **a,** Schematic of chronological aging in *S. cerevisiae*, illustrating the current model proposing impaired ribosome quality control (RQC) during aging.^16^ **b,** Quantification of pause-site and disome site frequencies across chronological aging time points. **c,** Violin plot showing increased elongation pausing during aging for pause sites detected across all aging time points. Center lines indicate the median, box boundaries indicate the interquartile range (IQR), and whiskers extend to 1.5 IQR. Pairwise comparisons across time points were performed using the Wilcoxon test (*n*=6,936): Pairwise comparisons were performed using the Wilcoxon test: day 0 *versus* day 2, \*\*\*\**p*= 1.61×10⁻⁶¹; day 0 *versus* day 3, \*\**p*= 1.68×10⁻⁵³; and day 2 *versus* day 3, ns, *p*= 0.247. **d,** Violin plot showing the effect of aging on co-translational ubiquitination (CTU). Center lines indicate the median, box boundaries indicate the IQR, and whiskers extend to 1.5 IQR. Statistical significance was assessed using the Wilcoxon test: \*\*\*\**p*= 1.4×10^-10^ (day 0, *n*=1,420; day 3, *n*=78). **e,** Gene ontology (GO) analysis of molecular function (MF) and cellular component (CC) terms for day 0 and day 3. **f,** Pearson correlation of amino acid enrichment at ribosomal E, P, and A sites for all disome sites on day 0 and day 3. Basic amino acids (lysine and arginine) are highlighted. **g,** Same as in **f**, but for codon optimality. *p*-values for differences between day 0 and day 3 were calculated by ANOVA. **h,** Heatmap of shared disome clusters undergoing RQC in at least one of the two time points. Color indicates the maximum log_2_(fold change) for each disome cluster, and clusters are hierarchically ordered based on adjusted *p*-values. Major clusters are highlighted. **i,** Metagene plot of monosome profiling data showing translation in the vicinity of day 0 ubiquitinated disomes and their corresponding non-ubiquitinated day 3 counterparts (*left, n*=22), and conversely day 3 ubiquitinated disomes and their corresponding non-ubiquitinated day 0 counterparts (*right, n*=10). Only genes with ≥200 normalized reads in both conditions within the analysis window were considered. Solid line represents the mean fraction of reads and shaded area the 95% CI. **j,** Volcano plot of PAN-Ub enrichment in ribosome-associated nascent chain (RNC) proteins on day 0 and day 3. Enrichment was calculated by comparing PAN-Ub pulldown samples with mock controls. Proteins associated with RQC, the ubiquitin-proteasome system, and ribosomal proteins are highlighted. AA, amino acid; CC, cellular component; GO, gene ontology; IQR, interquartile range; MF, molecular function; ORF, open reading frame; UPS: ubiquitin-proteasome system. n.s., *p*> 0.05; \**p*≤ 0.05; \*\**p*≤ 0.01; \*\*\**p*≤ 0.005; \*\*\*\**p*≤ 0.001.

### UbSeRP reveals endogenous RQC substrates

The RQC pathway has largely been defined using reporter systems containing ribosome stalling sequences and translation elongation inhibitors that result in prolonged ribosomal stalling and thereby promote disome accumulation *in vivo*.^1^ These tools enabled mechanistic dissection of RQC and delineated key steps by which cells recognize and clear deleterious disomes (**Fig. 2a**): (*i*) Hel2-dependent K63-linked polyubiquitination of the small ribosomal subunit of disomes; (*ii*) recruitment of the RQC trigger (RQT) complex for ribosome splitting; and (*iii*) engagement of Ltn1 on the 60S-nascent chain intermediate to drive extrusion and K48-linked ubiquitination of the nascent polypeptide followed by proteasomal decay.^12^

Although disomes are often considered a proteostatic burden,^12^ recent studies suggest that only a subset may engage in RQC, whereas others may be beneficial for translation elongation and may resolve spontaneously known as “ribosome cooperativity” (**Fig. 2a**).^24,25^ Thus, transcriptome-wide disome profiling alone may not specifically report on RQC,^14,15,26,27^ and in the absence of translatome-wide estimates, the prevalence of RQC-relevant disomes remains unclear. We therefore used UbSeRP to directly capture ubiquitinated disomes under endogenous conditions.

In total, we identified 36,466 disome sites across 2,645 genes. 6,938 sites in 937 genes were PAN-Ub-enriched, which included 2,162 sites in 443 genes that were also significantly enriched for K63-linked ubiquitin (**Fig. 2b**). Thus, ubiquitinated disomes comprise only a minority of the total disome population and frequently arise in transcripts associated with housekeeping functions, including metabolism and translation (**Supplementary Fig. 4a**). Importantly, across PAN-Ub-positive disome sites, PAN- and K63-Ub effect sizes were strongly correlated (**Supplementary Fig. 4b**), whereas K48-linked ubiquitin neither showed comparable enrichment nor correlated with PAN-Ub (**Fig. 2b**). This is likely due to the fact that Ltn1-mediated K48-ubiquitination occurs on split 60S ribosomal subunits, which have already released the transcript (**Fig. 2a**).^28^

In addition to recovering di-CGA codon motifs and premature termination-containing sites reminiscent of those in the known RQC substrates *sdd1* and *oaz1*,^4,5^ UbSeRP identified many additional sites with previously unknown sequence features, suggesting that endogenous RQC substrates are more diverse than appreciated (**Supplementary Fig. 4c**). To further confirm them as RQC sites, we assessed whether the identified sequences show the expected loss of ubiquitinated disomes in *hel2*Δ and stabilization in *slh1Δ*, as anticipated for RQC substrates (**Fig. 2c**). For this, we applied UbSeRP to *hel2*Δ and *slh1*Δ strains. We found that overall total disome levels were not severely altered in *hel2*Δ or *slh1*Δ genetic backgrounds (**Fig. 2d**, **Supplementary Fig. 4d**). Consistent with ubiquitinated disomes referring to RQC substrates, *hel2*Δ abolished ubiquitinated disomes, whereas *slh1*Δ increased and stabilized PAN- and K63-polyubiquitinated disomes relative to wildtype (**Fig. 2e**, **Supplementary Fig. 4e**). We also compared the data with a published massively parallel reporter assay of di-codon repeat sequences^13^ that measured mRNA stability in wildtype and *hel2*11 conditions.^29^ Our ubiquitinated disome sequences were enriched for sequence motifs identified in the massively parallel reporter assay, supporting that the majority of disomes signatures indeed resemble Hel2-sensitive sequences (**Supplementary Fig. 4f,g**).

We next asked whether ubiquitinated disome sites display translation defects due to defective RQC. To this end, we used monosome profiling data as a proxy for ribosomal occupancy and performed a metagene analysis centered on non-ubiquitinated and ubiquitinated disomes (**Fig. 2f**, **Supplementary Fig. 4h**). Reassuringly, ribosomal occupancy profiles around non-ubiquitinated disome sites did not differ across genetic backgrounds. In stark contrast, ubiquitinated disomes showed a broader ribosomal occupancy peak at the putative RQC site and pronounced differences across genetic backgrounds. In *slh1*Δ cells, ribosomes accumulated upstream of the putative RQC sites, implying translational defects, even forming stable trisomes. Downstream of these RQC sites, the ribosomal occupancy dropped markedly in *slh1*Δ, indicating that unresolved ubiquitinated collisions result in a translational roadblock. In contrast, this signature was not observed in *hel2*Δ and wildtype cells. This suggests that, in the absence of Hel2, collided ribosomes may resume translation spontaneously as suggested by reporter assays,^6^ whereas they are efficiently cleared by RQC in wildtype cells.

### Systematic identification of RQC determinants by UbSeRP

Our analysis indicated that defects in translation elongation were associated with two of the three ubiquitin signatures (**Fig. 1b**), namely clusters 2 (disomes) and 3 (pause sites). We therefore compared the molecular signatures of disome and pause site ubiquitin-dependent quality control. About 20% of disome sites were PAN-ubiquitinated and displayed strong K63-linked ubiquitin enrichment (**Figs. 2b** and **3a**). In contrast, only 1% of elongation pause sites were PAN-ubiquitinated and showed barely detectable K63-linked signal (**Fig. 3a**, **Supplementary Fig. 5a**). Moreover, even at the level of PAN-ubiquitin, the effect sizes differed significantly between pause and disome sites (**Supplementary Fig. 5b**), pointing to distinct modes of ubiquitin-dependent quality control. Together, our observations suggest that only a subset of pause sites and disomes lead to ubiquitination and RQC, whereas others may instead serve alternative roles in co-translational proteostasis. Therefore, we utilized our UbSeRP dataset to dissect the molecular signatures of ubiquitin-dependent quality control.

To define the determinants of these processes, we examined amino acid enrichment at the exit (E), peptidyl-transfer (P) and aminoacyl (A) sites, codon optimality, and biophysical properties of the nascent chain for paused ribosomes and disomes (**Fig. 5b**). We detected no striking differences in amino acid bias between total and PAN-Ub populations for either disome or pause sites (**Supplementary Fig. 5c,d**). The enriched arginine, lysine, glycine, and tryptophan across the E, P and A sites were frequently associated with optimal codons for total and PAN-Ub disome sites. We additionally found strong enrichment of stop codons at the A site of disomes. In contrast, pause sites more often encoded non-optimal codons, particularly at the A site, reminiscent of the elongation pause quality control pathway reported for the Ccr4-Not complex^30^ (**Fig. 3c**, **Supplementary Fig. 5f-h**). We did not detect significant differences in hydrophobicity between total and PAN-Ub disome or pause sites (**Fig. 3d, Supplementary Fig. 5i**). Together, these analyses indicate that neither amino acid identity, codon optimality, nor hydrophobicity alone is sufficient to explain the observed differences in ubiquitination of disome and pause sites.

We next examined additional biophysical features of the nascent chain that have been linked to translational stalling,^5,31,32^ but have not been resolved in the context of ubiquitin-dependent quality control previously. The isoelectric point (pI), which reflects charge distribution, was elevated at disome sites and was further increased in PAN-Ub disomes (**Fig. 3e**), whereas pause sites were associated with a reduced pI (**Supplementary Fig. 5i**), indicating that electrostatic interactions have differential effects on RQC pathways. PAN-Ub disomes and pause sites also showed increased nascent chain bulkiness relative to their corresponding total populations (**Fig. 3f; Supplementary Fig. 5i**), consistent with steric hindrance in the exit tunnel potentially contributing to elongation defects. For disomes, this effect peaked ∼10–15 codons upstream of the A site (**Fig. 3f**), matching the approximate position of exit tunnel constriction. Together, our systematic disome-wide analysis showed that elongation stalls associated with ubiquitin-dependent quality control are characterized by more positively charged nascent chains and greater bulkiness than their non-ubiquitinated counterparts. This is in agreement with a previous structural dissection of the *sdd1* RQC site.^5^

Beyond sequence and biophysical features, we furthermore considered if the structural conformation of disomes influences RQC. Disome footprints displayed a bimodal length distribution (**Supplementary Fig. 1c**), corresponding to proximal (59-61 nt) and distal (63-65 nt) disomes (**Fig. 3g**).^14,15,26,27^ Comparison of the effect size between PAN- and K63-Ub disomes revealed that the fold change of proximal disomes was significantly higher with respect to the distal disomes (**Fig. 3h**). Interestingly, ubiquitinated disomes in the *slh1*11 background only contained proximal disomes suggesting that proximal disomes may be the substrate of RQC (**Supplementary Fig. 5j**). Our findings imply that the disome geometry may affect ubiquitination and may resemble an important structural intermediate in RQC.

### RQC intersects with co-translational mRNA quality control and protein folding

RQC can result in the activation of mRNA surveillance pathways and ultimately mRNA degradation (**Fig. 3i**).^33^ We therefore stratified RQC-associated transcripts, rather than disome-associated transcripts, to obtain a more resolved view of their association with mRNA surveillance and degradation. Employing the codon protection index, a metric reporting on co-translational mRNA decay, derived from a previously published 5PSeq dataset,^34^ we correlated mRNA decay with disome and ubiquitination frequencies. We found that transcripts with lower disome frequencies were more susceptible to co-translational decay (**Fig. 3j**). Strikingly, decay-prone transcripts were enriched with high ubiquitination frequency (**Fig. 3j**), suggesting that co-translational mRNA decay is not strictly determined by disome frequency, but involves disome ubiquitination.

In addition to mRNA decay, disomes and RQC have been implicated in protein folding, including the biogenesis of TMDs and protein complexes.^14,21,27,35^ We observed a significant enrichment of RQC near TMDs as well as previously annotated Ssb1/Ssb2 and TRiC binding sites (**Fig. 3k**), supporting a role of RQC in co-translational protein folding. In contrast, protein dimerization domains associated with co-translational assembly (co-co domains)^21^ were enriched for non-ubiquitinated disomes (**Fig. 3k**). Together, our findings suggest that disomes may have different roles for co-translational proteostasis.

### A model to predict RQC engagement

Given the diversity of potential convoluted causes that distinguish RQC-engaged from non-engaged disomes, we trained a random forest classifier using ubiquitination status, sequence bias, biophysical and structural features, and chaperone engagement as input variables (**Fig. 3l**). The resulting model discriminated ubiquitinated disomes from total disomes with substantial predictive power, achieving an area under the receiver operating characteristic (ROC) curve of 0.86 and a precision-recall curve of 0.64 (**Fig. 3m**, **Supplementary Fig. 5k)**. Notably, the classification was not driven by a single feature but a combination of factors. Strong contributions arose from nascent chain and mRNA intrinsic features, including the pI, codon optimality, amino acid bulkiness, and hydrophobicity (**Fig. 3n**). Intriguingly, features such as chaperone engagement or TMD contributed comparatively little to the predictability of RQC.

### Co-translational nascent chain ubiquitination resembles a distinct quality control pathway

Unlike CTU_RQC_, CTU_NC_ is less well understood, where substrates, their modification sites and E3 ligases remain largely unknown (**Fig. 4a**). As co-translational misfolding is increasingly recognized,^37^ defining quality control on nascent chains is essential. Using UbSeRP, we identified 138 PAN-, 5 K48-, and 3 K63-ubiquitinated nascent proteins, with overlap between PAN and each linkage type-specific set, but none between K48 and K63 (**Fig. 4b-d**, **Supplementary Fig. 6a,b**).

To reveal features associated with CTU_NC_, we tested whether onset sites localize near degron motifs. Indeed, degron sequences accumulated near the onset of CTU_NC_ of the nascent chain, whereas RQC sites were not enriched, suggesting that E3 ligases can target nascent polypeptides independently of RQC (**Fig. 4e**). Interestingly, the onset of co-translational ubiquitination only coincided with previously identified post-translational ubiquitination sites in a few cases,^38^ raising the possibility that these sites are targeted specifically during co-translational protein biogenesis. Using co-translational chaperone-binding sites as proxies for misfolding-prone nascent chain regions, we found that CTU_NC_ onsets did not overlap with these sites. Instead, we found that CTU_NC_ onsets frequently occurred in 30 amino acid proximity to or within structured protein domains (**Fig. 4f**). In particular, many CTU_NC_ onsets coincided with nucleic acid binding sites and protein-protein interaction domains, such as coiled-coils **(Fig. 4g)**, which have been recently described to assemble co-translationally.^21^ Structural mapping of the predicted degrons onto protein structures indicated that degrons encoded in structured domains may be accessible in the isolated subunit but may become buried within protein-protein interfaces upon assembly, suggesting that protein complex formation may mask ubiquitination sites (**Supplementary Fig. 6c**).

### Aging affects translation and reveals novel insights into the dynamics of RQC

Proteostasis declines during aging and is associated with reduced translational capacity and impaired ribosome function.^42^ Notably, pronounced elongation defects have been proposed to drive ribosome collisions and overwhelm RQC (**Fig. 5a**).^16,17^ However, direct support for this model is largely lacking, in part because data linking ribosome collisions to ubiquitin signaling was missing.

To define how aging remodels translation and translation-coupled quality control, we chronologically aged *S. cerevisiae* and analyzed cells on days 0, 2 and 3 using UbSeRP and quantitative proteomics. Our translatome and proteome data recapitulated established hallmarks of aging,^43^ including a reduced abundance of factors involved in ribosome biogenesis, translation and protein folding, and an increase in stress-responsive components linked to ubiquitination, proteolysis, chaperone function and the integrated stress response (**Supplementary Fig. 7a-c**). Mass spectrometry of purified RNCs on day 3 compared to day 0 showed a reduced association of co-translational factors, including chaperones and SRP, and an increased association of ubiquitin-proteasome components (**Supplementary Fig. 7d**).

We next examined whether this aging-associated remodeling was accompanied by elongation defects and found that puromycin incorporation into nascent chains, which we used as a proxy of translation elongation, declined with age (**Supplementary Fig. 7e**). Although pausing and disome formation were overall reduced in aged cells, conserved pause sites showed increased pause scores (**Fig. 5b,c**), consistent with previous reports.^16^ Interestingly, age-emergent disomes preferentially mapped to a distinct set of transcripts with increased translation during aging (**Supplementary Fig. 7f**).

Thus, we asked how aging affects CTU, particularly RQC, and found that aging reduced the effect size of PAN-Ub enrichment, while the ratio of PAN-Ub sites to non-ubiquitinated disomes ranged between ∼2.3% (day 3) - ∼3.0% (day 0) (**Fig. 5d**). Additionally, we found that RQC occurred more frequently in genes associated with mitochondrial protein complexes and translation elongation factors (**Fig. 5e, Supplementary Fig. 7f**). Many of these disomes more frequently formed on non-optimal codons with age, which is in line with altered cellular energy metabolism in aging which regulates mRNA stability^44^ (**Fig. 5f,g**). We also detected disomes sites that were conserved during aging, but which were ubiquitinated only at one timepoint and not the other, revealing a context dependence of disome recognition and ubiquitination (**Fig. 5h, Supplementary Fig. 7g,h**). To assess whether translation at the differentially ubiquitinated disomes is altered, we performed a metagene analysis. Our data revealed that translation efficiency after these disome sites was only mildly affected in ubiquitinated disomes at day 3 compared to the non-ubiquitinated counterparts from day 0 (**Fig. 5i, Supplementary Fig. 7i**).

To validate our UbSeRP observations, we performed quantitative proteomics of PAN-Ub enriched RNCs and found that they were more strongly associated with canonical RQC factors and the Ccr4-Not complex in young cells, whereas aged PAN-Ub RNCs were enriched for proteasome-linked factors, including the Ufd1-Npl4-Cdc48 complex, K48-ubiquitin readers, proteasomal lid components, and Tom1 (**Fig. 5j**). Together, these findings indicate that aging does not necessarily overwhelm RQC. Rather, age-associated RQC events may arise at distinct sites across different genes. Furthermore, aging may result in the engagement of proteasome-linked degradation.

## Discussion

Protein biogenesis requires tight coordination of mRNA decoding by the ribosome, protein folding, and co-translational assembly.^8,45^ However, errors in translation can stall ribosomes, causing them to collide and form disomes, as well as generating aberrant nascent chains that threaten proteome integrity.^1^ Although ubiquitin-dependent quality control can mitigate this risk for proteostasis,^2,3^ it has remained unclear which translational states are surveilled and how cells differentiate those states from productive translation.

By interrogating ubiquitination of ribosome-nascent chain complexes, UbSeRP enables codon-resolved mapping of co-translational ubiquitination (CTU) to address these questions. To achieve this, we rely on TUBEs that have distinct intrinsic affinities for specific linkage types and chain architectures.^18^ Unlike reporter- or ribosome profiling-based approaches, such as disome profiling in RQC-deficient genetic backgrounds^15^ or selective ribosome profiling of individual E3 ligases,^30,46,47^ UbSeRP directly quantifies ubiquitination, distinguishes ribosomal from nascent chain modification, and provides an unbiased, translatome-wide view of CTU (**Fig. 1a**). We show that subsequent integration of the multimodal UbSeRP datasets resolve CTU for distinct translational states, separating ribosome pausing and disome formation from nascent chain ubiquitination (**Fig. 1b-e**).

Using this approach, we uncover widespread nascent chain ubiquitination independent of RQC, revealing an unrecognized layer of CTU. RQC has been proposed to monitor deleterious co-translational assembly, particularly at elongation stalls near assembly onset sites that can lead to disome formation.^45^ However, disomes at these sites are frequently not ubiquitinated and instead localize to dimerization domains implicated in co-translational assembly (**Fig. 3k**),^21^ suggesting assembly-associated pausing rather than quality control engagement. In contrast, CTU_NC_ ubiquitination occurs within similar interaction domains but independently of disomes, indicating that nascent chains can be directly ubiquitinated (**Fig. 4e-g**), which may be a necessity to regulate orphaned nascent subunits that can adopt assembly incompetent folding states once released from the ribosome.^48^ Notably, co-translational ubiquitination sites do not coincide with post-translational ubiquitination marks (**Fig. 4e**), indicating that these events are temporally restricted to early biogenesis, before the ubiquitination sites become buried within structural interfaces upon assembly (**Supplementary Fig. 6c**) and may interfere with mature protein function.

Interestingly, CTU_NC_ displays a distinct linkage composition, with limited K48 enrichment (**Fig. 4b, Supplementary Fig. 6a**), despite the association of K48-linked polyubiquitin chains with CTU.^2^ Notably, many K48-linked polyubiquintated nascent chains may be short-lived and highly substoichiometric. Therefore, we may fail to detect them robustly. In contrast, we find some nascent chains with K63-linked ubiquitination, which may hint at regulatory roles of CTU_NC_ in protein biogenesis. Consistently, many UbSeRP-defined domains overlap with conserved ubiquitination sites associated with regulatory, rather than degradative, functions, suggesting that these are not targets of K48-linked ubiquitination.^49^ Together, CTU_NC_ may regulate protein complex assembly.

Furthermore, our data challenge the prevailing view that disomes are inherently deleterious signals that trigger RQC. We find that disomes are highly abundant across the yeast translatome, yet only a minority is ubiquitinated (**Fig. 2b**). Consistently, non-ubiquitinated disomes do not impair translation in RQC-deficient strains, indicating that most disomes are compatible with protein synthesis (**Fig. 2f**).

These findings strengthen the emerging model of “ribosome cooperativity”^24^ in which the majority of disomes facilitate productive translation by alleviating ribosome pausing at problematic sequences. Ribosome cooperativity resolves the apparent paradox whereby highly expressed proteins such as ribosomal proteins and histones, whose sequence are reminiscent of the poly-basic reporter constructs used to determine the mechanism of RQC,^1^ are efficiently synthesized despite increased disome frequencies. While such cooperativity via short-lived disomes allows most ribosomes to support productive elongation across difficult sequences, it comes at the cost of a subset of deleterious disomes that stall and require RQC. Thus, balancing translational demand with quality control enables high protein output while preserving proteome integrity, with RQC acting as a constitutive housekeeping pathway in non-stressed cells.

To further utilize UbSeRP, we examined chronological aging, where RQC impairment has been proposed due to an observation of increased ribosome pausing in yeast, killifish and nematodes.^16,17^ Yet, disomes and RQC activity was not directly measured on endogenous substrates. Our experiments revealed two major insights into the adaptability of co-translational ubiquitination. The RQC regulatory capacity exhibits a striking adaptability. Many disome sites ubiquitinated in young cells were not detected as ubiquitinated in aged cells, and vice versa (**Fig. 5h**). In particular, changes in tRNA availability and the metabolic state of a cell associated with aging^17,44^ may be involved in determining which disomes become deleterious. Overall, our analysis of the UbSeRP datasets demonstrates that substrate selection for RQC is rather intricate and is likely regulated at multiple levels. Therefore, systematic application of UbSeRP across diverse cellular states will be essential to define the underlying decision logic of RQC engagement.

Although aging is associated with an accumulation of misfolded proteins,^9^ RQC remains functional, as shared disome sites in young and aged cells do not exhibit translational defects (**Fig. 5i**) comparable to those observed in RQC-deficient strains (**Fig. 2f**). Instead, we find that, while elongation pausing is increased during aging as reported previously (**Fig. 5c**),^16,17^ this is not accompanied by elevated disome abundance (**Fig. 5b**). Rather, overall disome levels decrease. One possible explanation is that age-associated defects in translation elongation reflect reduced ribosome cooperativity, such that pausing increasingly occurs, for example on polybasic stretches that were alleviated by cooperative disomes in young cells. Rather than an age-related overload or collapse of RQC at disomes as previously hypothesized, our data point to a rerouting of aberrant translational states towards proteasome-mediated clearance (**Fig. 5j**), as indicated by the increased association of proteasomal components and ubiquitin shuttle proteins with RNCs.

Together, UbSeRP and the accompanying multimodal dataset provide a foundational and rich resource for a broader community. By resolving endogenous substrates for RQC, this work shifts the investigation of RQC from synthetic reporters to native targets and enables mechanistic dissection through biochemical and structural approaches. Our insights into RQC and the UbSeRP framework may inform rational mRNA vaccine design, where limiting unwanted RQC-mediated mRNA destabilization is critical,^50^ and enable mechanistic studies of e.g. RQC dysregulation in neurodegenerative diseases such as Huntington’s disease.^51^ Finally, some significantly elevated ubiquitination sites within the translatome remain unassigned to the three quality control pathways described here (CTU_NC_ and CTU_RQC_ on disomes and paused ribosomes), suggesting additional quality control mechanisms. While their nature remains unclear, this dataset provides a basis for hypothesis generation and the discovery of additional quality control pathways. Overall, our findings establish CTU as a dynamic, context-dependent surveillance layer whose complexity extends beyond currently recognized pathways.

## Data availability

The data will be made available upon publication.

## Code availability

The code will be made available upon publication.

## Author contributions

M.S. conceived the project, designed and performed experiments, analyzed data, and wrote the manuscript. S.L. analyzed data and wrote the manuscript. K.G. performed experiments, analyzed data, and wrote the manuscript. X.H.A. performed analysis and wrote the manuscript. I.K. performed experiments, analyzed data, and wrote the manuscript. Z.K., A.B., D.K.W. and B.K. performed experiments. J.P. and J.W. analyzed data. D.N., J.G., J.D.L., V.B., M.M.S., and J.F. supervised the project. M.B. and L.M.S. conceived and supervised the project and wrote the manuscript.

## Acknowledgement

We thank Stefanie Böhm, Carlos Alfonso Gonzalez, Mira Burtscher, Martin Garrido Rodriguez, Wolfgang Huber, Melanie Krause, Marta Seczynska, and Margret Wangeline for insightful discussion of the project. We also want to thank Angela Chu from the Stanford Genome Technology Center for providing the *S. cerevisiae* BY4741 *hel211* and *slh111* strains. We also like to thank Erin Schuman and Claudia Fusco at the Max Planck Institute of Brain Research for polysome profiling. Furthermore, we would like to thank Sonja Welch and the electron microscopy facility at the Max Planck Institute of Biophysics. The authors thank Charles Girardot from the EMBL Multimodal Open Data Integration Support. M.S. received funding through the Life Science Alliance Bridging Excellence Fellowship. X.H.A. was supported by a Humboldt Research Fellowship. J.D.L. acknowledges funding by the Max Planck Society. V.B., M.M.S., and L.M.S. acknowledge their funding by EMBL. The project was funded by the Max Planck Society, and the European Research Council granted to M.B. (724349-Complex assembly).

## Declaration of generative AI and AI-assisted technologies in the writing process

During the preparation of this work, the authors used ChatGPT (OpenAI, GPT-5.4) in order to generate and improve code and revise text for the manuscript. After using ChatGPT, the authors reviewed and edited the content as needed and take full responsibility for the content of the published article.

## Methods

### Yeast strains and growth conditions

All experiments were performed in *Saccharomyces cerevisiae* BY4741. BY4741 *slh1*11 and *hel2*11 were deleted using a kanMX4 cassette and were obtained as part of the genome-wide deletion library from the Stanford Genome Technology Center.^54^ For UbSeRP in YPD, *S. cerevisiae* was plated on YPD two days before the experiment. Then, single colonies were picked and grown overnight at 30°C and 180 rpm. On the next day, OD600 was measured. 4 x 400 mL (total: 1.6 L) in 2 L flasks per biological replicate was set to an OD600 of 0.035 and grown to an OD600 of 0.5-0.6 at 30°C, 160 rpm. Subsequently, cells were harvested by rapid filtration onto a nitrocellulose membrane (0.45 µm, Biorad) and flash-frozen in liquid nitrogen.

For the aging experiments, *S. cerevisiae* was grown in synthetic complete media containing five-fold access for auxotrophic nutrients (SC+ media) as described in Stein et al.^16^ Therefore, the following concentrations of uracil (0.875 g/L), leucine (1.75 g/L), methionine (0.875 g/L), and histidine (0.875 g/L) were used. For each aging experiment, yeast was first grown on YPD two days before the experiment. Then, single colonies were inoculated in SC+ media overnight at 30°C, 180 rpm. In analogy to the experiments in YPD, OD600 was determined, and 6 x 400 mL (total: 2.4 L) in 2 L flasks per biological replicate was set to an OD600 of 0.035 and grown at 30°C, 160 rpm. 800 mL of cells were harvested at an OD600 of 0.5-0.6 (day 0, OD600 ∼500-600). For day 2 (48 hrs post-inoculation) and day 3 (72 hrs post-inoculation), 100 mL of cultures were harvested, approximating to an OD600 of 500-600. The harvest and freezing were performed as stated above.

To further ensure the reproducibility of the aging experiments, all colonies subjected to the aging experiments were additionally grown on YP-glycerol. Successful growth of cells on YP-glycerol suggests that cells are capable of respiration, as glycerol corresponds to a non-fermentable carbon source. Additionally, we confirmed cell viability at each time point employing the Yeast Viability Kit I (Logos Biosystems) according to the manufacturer’s instructions. Cell viability was read out using a LUNA-FLTM automated cell counter. The cell viability usually ranged from 90-98% suggesting high viability of the aged cultures.

### Ubiquitin selective ribosome profiling

The presented method is an adaptation of the previously published SeRP workflow.^52,55–57^. Frozen cells were supplemented with 2.4 mL of high salt lysis buffer (20 mM Hepes-KOH, pH 7.5, 500 mM KCl, 20 mM MgCl_2_, 1 mM PMSF, 0.01 % IGEPAL, and cOMPLETE EDTA-free protease inhibitor (Roche), 0.1 mg/mL CHX (Sigma-Aldrich) or 0.01 mg/mL puromycin (Sigma-Aldrich)). Afterwards, cells were lysed using the CryoMill (Retsch) at 30 Hz for 2 min. After lysis, lysates were gently thawed at 4 °C and supplemented with an additional 2.4 mL of high salt lysis buffer containing 40 mM NEM and 40 µM phosphoramidon to inhibit deubiquitination and degradation. The lysate was cleared by centrifugation at 15,000 *g* at 4 °C for 3 min.

Next, the absorbance at 260 nm (A260) of a 1:100 dilution was measured by nanodrop to estimate the amount of RNase I. To convert the polysomes into mono- and disome, we supplemented the lysate with 100 U per A260. The RNase I digest was mixed by end-to-end rotation and incubated for 40 min at 4°C. Afterwards, the reaction was immediately quenched by adding 600 U of Superase•In.

To enrich for ribosome-nascent chain complexes (RNCs), lysates were loaded onto 25 % sucrose cushions (20 mM Hepes-KOH pH 7.5, 140 mM KCl, 10 mM MgCl2, 25% w/v sucrose, 0.01% IGEPAL, 0.1 mg/mL CHX or 0.01 mg/mL puromycin, 1 tablet of cOMPLETE protease inhibitor per 50 mL, 20 mM NEM, 20 µM phosphoramidon). The RNCs were pelleted at 150,000 *g* and 4 °C for 2.5 hrs. Next, the sucrose cushion was removed from the RNC pellet and the pellets were resuspended in 1 mL of wash A (20 mM Hepes-KOH pH 7.5, 140 mM KCl, 10 mM MgCl2, 0.01% IGEPAL, 0.1 mg/mL CHX or 0.01 mg/mL puromycin, 1 tablet of cOMPLETE protease inhibitor per 50 mL, 20 mM NEM, 20 µM phosphoramidon). From the RNCs, 100 µg of RNA was used for the total translatome libraries of mono- and disomes. The remaining sample was split for each condition and added to 125 µL of pre-equilibrated streptactin resin (IBA). To enrich for the different ubiquitin linkages, following TUBEs were added: PAN-TUBE1 (biotin) (12.5 µg, LifeSensors, UM-0301-0200), K48-TUBE HF (biotin) (12.5 µg, LifeSensors, UM-0307), and K63-TUBE (biotin) (12.5 µg per experiment; LifeSensors, UM-304-0050). Beads were incubated for 90 min at 4°C while end-to-end mixing. Afterwards, beads were washed three times with wash A, each 1 min while end-to-end mixing, followed by two washes in wash B (20 mM Hepes-KOH pH 7.5, 140 mM KCl, 10 mM MgCl2, 0.05% IGEPAL, 10 % v/v glycerol, 0.1 mg/mL CHX or 0.01 mg/mL puromycin, 1 tablet of cOMPLETE protease inhibitor per 50 mL, 20 mM NEM, 20 µM phosphoramidon). The first wash was performed for 1 min, and the second wash for 4 min, while end-to-end mixing. After washing, beads were resuspended in 500 µL 10 mM Tris-HCl, pH 8.0, and 40 µL of 20 % SDS and gently inverted. Next, 750 µL of pre-warmed phenol-chloroform-isoamyl alcohol (PCI, 65°C) was supplemented, and mixed at 1400 rpm at 65°C for 5 min. Immediately after, Eppendorf tubes were placed on ice for 10 min and then centrifuged at 15,000 *g* for 10 min. Subsequently, the aqueous phase was transferred into a fresh Eppendorf tube, and the PCI extraction was repeated, but at room temperature. After the second extraction, the aqueous phase was mixed with 500 µL of diethyl ether, vortexed for 5 min, centrifuged for 10 min at 4°C, and the aqueous phase was isolated. Residual organic solvent was evaporated using a SpeedVac at room temperature for 30 min. The aqueous phase was then supplemented with 3 M NaOAc, pH 5.5, to obtain a final concentration of 0.3 M, 2.5 µL of Glycoblue (Invitrogen), and equal volumes of isopropanol relative to the volume of the aqueous phase. The precipitation reaction was vigorously vortexed and incubated at -80°C overnight. To precipitate the RNA, tubes were centrifuged at 15,000 *g* and 4°C for 90 min. The RNA pellet was then washed twice with 70 % ice-cold EtOH. Subsequently, pellets were dried in the SpeedVac at room temperature for 30 min and resuspended in 20 µL TE buffer.

The following adaptations were made for the aging experiments: (*i*) cells obtained from day 0 were lysed in 2.4 mL of high salt lysis buffer, while 4.8 mL of high salt buffer was used to lyse the cells obtained from day 2 and day 3. The lysis buffer was supplemented with 20 µM PB-619 (Sigma Aldrich), and 20 mM NEM. (*ii*) 75 µL of magnetic streptavidin (Thermo Scientific) beads with 25µg of PAN-TUBE1 (biotin) (LifeSensors, UM-0301-0200) were used.

The resuspended RNA was mixed with an equal volume of 2x RNA loading dye (Thermo Scientific). The RNA marker was prepared with 200 nM synthetic 5′FAM-labeled RNAs. For monosomes, we isolated footprints in a range of 26 to 34 nucleotides, while dimers were isolated from 54 to 68 nt. Sequences of the custom-made RNAs can be found in **Supplementary Table 2**.

**Supplementary Table 1:**
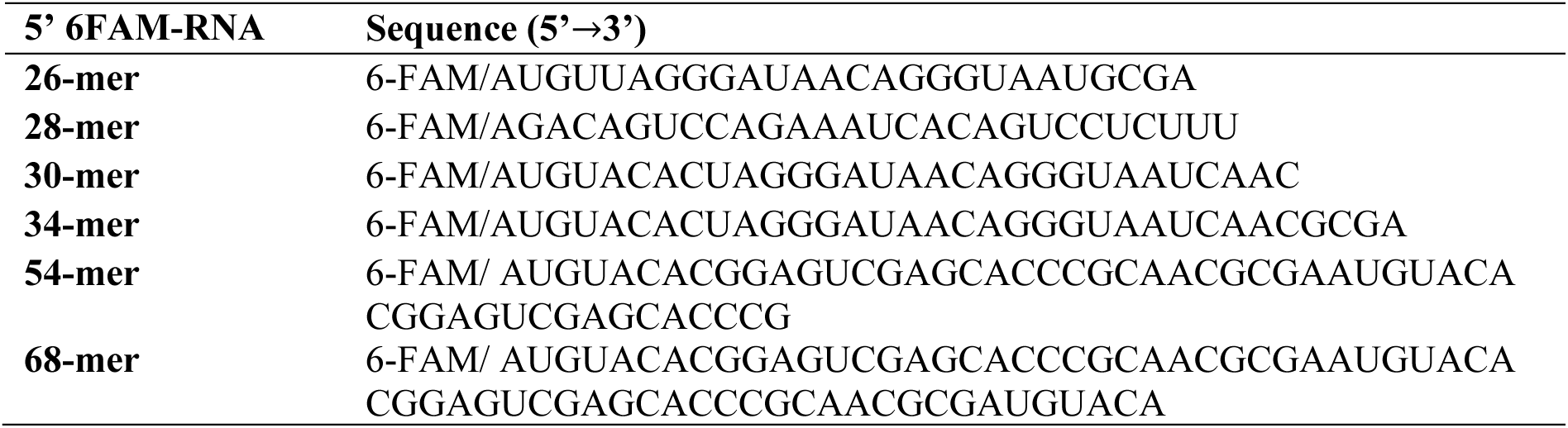
Sequences for ribosome profiling markers. 26-34-mers were used as a reference to excise monosomes as described in^55,56^. Disomes were obtained using the 54-68-mer marker. The sequences for the disome marker are from ^15^.

Both the sample and the marker were denatured at 80°C for 2 minutes and then placed on ice. A 15% denaturing PAGE (Carl Roth) was prepared, prewarmed for 1 hour at 16 W, and then loaded. The gels were run at 16 W for 3.5-4 hours until the bromophenol blue eluted from the gel. After running, the gels were stained with SybrGold (Invitrogen) and imaged using the Amersham Typhoon (GE Healthcare). The gel regions between 26 to 34 nt and 54 to 68 nt were excised and crushed. RNA was eluted in 500 µL of Tris-HCl pH 8.0 at 70°C for 10 minutes while shaking at 1,400 rpm. The eluate was separated from the gel pieces using a Spin-X cellulose acetate column with a 0.22 µm pore size (Corning). RNA was precipitated by adding 50 µL 3 M NaOAc pH 5.5, 2.5 µL Glycoblue co-precipitation agent (Invitrogen), and 500 µL isopropanol.

The purified RNA was initially dephosphorylated in 1x FastAP buffer containing 2 U FastAP (Thermo Scientific) and 20 U RiboLock (Invitrogen). This reaction was incubated at 37°C and 600 rpm for 15 minutes, followed by immediate heat inactivation at 75°C for 5 minutes. The 5′ ends were then phosphorylated using 20 U polynucleotide kinase (PNK; NEB) with 1 mM ATP (Thermo Scientific), 1x PNK buffer (NEB), and 20 U RiboLock. This reaction was incubated for 30 minutes at 37°C.

RNA quantity was determined using the Qubit RNA high sensitivity kit. For library preparation, inputs were standardized to 25 ng for monosomes and 100 ng for disomes. Disome inputs were rRNA depleted using yeast specific rRNA depletion probes following the riboPOOL ribosomal depletion kit (siTOOLS Biotech). Monosome libraries were prepared using the NEXTflex small RNA sequencing kit v3 and the disome libraries were prepared using the NEXTflex small RNA sequencing kit v4 as per manufacturer instructions. For *slh1*11 and *hel2*11 UbSeRP experiments, libraries were prepared using the NEBNext low-bias small RNA library preparation kit. Libraries were sequenced on an Illumina NextSeq2000 using a P3 kit 50 base pair single-end for monosomes and a P3 kit 50 base pair paired-end for disomes.

### Analysis of CTU and translation remodeling

100 mL of *S. cerevisiae* BY4741 was grown in SC+ for each time point as described above. Cells were harvested by centrifugation at 8000 *g* at 4°C for 2 min. The pellets were washed in 1000 µL PBS and transferred into Eppendorf tubes, and the centrifugation step was repeated to remove the PBS. Cells were lysed in 500 µL high salt ribosome profiling buffer (20 mM Hepes-KOH pH 7.5, 500 mM KCl, 10 mM MgCl2, 0.01% IGEPAL, 0.01 mg/mL puromycin, 1 tablet of cOMPLETE protease inhibitor per 50 mL, 20 mM NEM) in the CryoMill (2 min, 30 Hz).

The lysate was thawed and centrifuged at 15,000 *g* at 4°C for 3 min. 100 µL of the lysate was taken and subjected to BCA. The other 400 µL of the lysate was loaded onto 800 µL of sucrose cushion buffer (20 mM Hepes-KOH pH 7.5, 140 mM KCl, 10 mM MgCl2, 25% w/v sucrose, 0.01% IGEPAL, 0.01 mg/mL puromycin, 1 tablet of cOMPLETE protease inhibitor per 50 mL, 20 mM NEM) and centrifuged at 245,000 *g* at 4°C for 90 min. 100 µL of the top layer was taken as supernatant. The residual sucrose cushion was removed, and the RNC pellet was resuspended in 400 µL of wash A (20 mM Hepes-KOH pH 7.5, 140 mM KCl, 10 mM MgCl2, 0.01% IGEPAL, 0.01 mg/mL puromycin, 1 tablet of cOMPLETE protease inhibitor per 50 mL, 20 mM NEM). Both, supernatant and resuspended RNCs, were subjected to BCA to determine protein concentration. Protein concentrations of lysate, supernatant, and RNCs were adjusted to 10 µg/µL and mixed with 4 x NuPAGE loading dye. 10 µL of each fraction was loaded per lane.

### Sample preparation of ribosomes for proteomics analysis

*S. cerevisiae* BY4741 were grown as described above using the SC+ media. Ribosomes were purified from 400 mL at day 0 and 100 mL at day 3.

In brief, cells were lysed in 1.25 mL of lysis buffer (20 mM Hepes-KOH pH 7.5, 140 mM KCl, 10 mM MgCl2, 25% w/v sucrose, 0.01% IGEPAL, 0.1 mg/mL CHX, 1 tablet of cOMPLETE protease inhibitor per 50 mL) in a CryoMill at 30 Hz for 2 min. After gently thawing, lysates were mixed with 1.25 mL of ribosome profiling buffer containing 40 mM NEM, 80 µM PR-619 to inhibit deubiquitination. Next, lysates were cleared at 15,000 *g* for 3 min at 4°C. Then, the cleared lysates were loaded on 25 % sucrose cushions (20 mM Hepes-KOH pH 7.5, 140 mM KCl, 10 mM MgCl2, 25% w/v sucrose, 0.01% IGEPAL, 0.1 mg/mL CHX, 1 tablet of cOMPLETE protease inhibitor per 50 mL, 20 mM NEM, 40 µM PR-619). The sucrose cushions were run for 150,000 *g* and 4°C for 2.5 hrs.

After RNC resuspension in 750 µL of wash A (20 mM Hepes-KOH pH 7.5, 140 mM KCl, 10 mM MgCl2, 0.01% IGEPAL, 0.1 mg/mL CHX, 1 tablet of cOMPLETE protease inhibitor per 50 mL, 20 mM NEM, 20 µM phosphoramidon), the protein concentration was determined using the BCA assay (Pierce). 30 µg of total RNC was subjected to MS sample preparation to obtain the total RNC proteome, while the residual sample was split into two 350 µL fractions and loaded on 45 µL pre-equilibrated PAN-Ub (LifeSensors, Cat No. UM-0501M-1000) and mock pulldown beads (LifeSensors, Cat No. UM-0500M-1000). Before loading on beads, purified polysomes were digested with 10 U of RNase I per absorbance unit for 10 min at 4°C and immediately quenched using 20 U Superase•In. RNCs were incubated on the beads and mixed by end-to-end mixing for 1.5 hrs at 4°C. After incubation, the beads were washed four times with 1 mL of wash A. The last wash was removed, and beads were boiled at 72 °C in 1x NuPAGE loading buffer. 5 µL of the beads were subjected to Western Blot analysis.

Briefly, RNC samples, PAN-Ubiquitin enriched RNCs, and controls were adjusted to a final SDS concentration of 5% (w/v). Cysteines were reduced by the addition of 20 mM DTT, followed by alkylation with 50 mM iodoacetamide. Samples were acidified with 1.2% phosphoric acid and cleared at 20000 *g* for 10 min at room temperature. The supernatant was diluted using TEAB-buffered Methanol at a ratio of 1:7.

Proteins were trapped on S-Trap micro cartridges (Protifi) by centrifugation for 30 sec at 4000 *g* and washed four times with TEAB-buffered Methanol. 1 µg of Trypsin (Promega, Sequencing Grade Modified Trypsin) in 60 µL 50 mM TEAB was applied atop the cartridge. Digestion was allowed to proceed for approximately 16 hrs at room temperature in a humidified chamber. Peptides were eluted by centrifugation at 4000 *g* for 60 sec once with 40 µL 50 mM TEAB and twice with 40 µL 0.2 % formic acid. Pooled eluates were desalted using two disks of Empore 3M C18 material as described in Rappsilber et al.^58^ and dried in a SpeedVac.

Dried peptides were solubilized in buffer A (5% acetonitrile, supplemented with 0.1 % FA) and subjected to LC-MS/MS analysis on a timsTOF HT instrument (Bruker) with a nanoElute (Bruker) system. Peptides were separated on a heated C18 column (60°C, PepSep 150 mm x150 µm ID, 1.5 µm particle size (Bruker)) at a flow rate of 800 nL/min using the following gradient: 2 to 38% B in 21 min, 38 to 95 % B in 0.5 min and constant 90 % B for 3.5 min (buffer A: 0.1 % FA in water; buffer B: 80 % acetonitrile, 0.1 % FA in water).

Eluting peptides were ionized using a captive spray ion-source and analyzed in DIA-PASEF mode with a cycle time of 0.96 s and 25 Da PASEF windows. Spectra were acquired over the mass range from 100-1,700 m/z and a mobility window from 0.85-1.3 Vs/cm^2^.

Peptide identification and label-free quantification were performed in DIA-NN 1.8.2 beta (nascent chain fractions and pulldowns)^59^ against the *S. cerevisiae* Uniprot reference proteome (UP000002311, accessed 15.11.23).

The database search space was restricted to tryptic peptides with a length of 7-30 amino acids, 1-4 charges, and a m/z range of 300-1,800, allowing for up to one missed cleavage. Carbamidomethylation of Cysteine was set as a fixed modification and K-GG was set as variable modification. Automated mass accuracy optimization was selected, Match-Between-Runs was enabled and data normalization was disabled. The results were FDR-filtered (FDR < 0.01).

The mass spectrometry proteomics data will be uploaded to the ProteomeXchange Consortium (http://proteomecentral.proteomexchange.org) via the PRIDE partner repository.^60^

The resulting DIA-NN output results were further analyzed using the MS-DAP pipeline (V1.0.6).^61^ Peptides were filtered and normalized for each pairwise comparison (“contrast filtering”) with the following settings: fraction_detect = 0.75; fraction_quant = 0.75; min_peptide_per_prot = 3; norm_algorithm = ‘vsn’; rollup_algorithm = ‘maxlfq’. The implemented msqrob algorithm was used for differential expression analysis with an FDR threshold of q < 0.05 and a log_2_fold change threshold of log_2_(1.5) for the analysis of IPs and log2(fold change) ≥ 0.75 to detect differentially abundant proteins in the RNCs.

### Western Blot analysis

Efficiency of ubiquitin enrichment (**Supplementary Fig. 1b**) and age-associated defects in translation (**Supplementary Fig. 7e**) were assessed using Western Blotting. In brief, protein samples were mixed with 4 x NuPAGE loading dye containing 5 % beta-mercaptoethanol to obtain a final concentration of 1 x NuPAGE loading dye. Protein was denatured at 72°C for 5 min before loading on a NuPAGE Bis-Tris gel. Unless otherwise stated, 4 µL of lysates and RNCs, and 10 µL of the bead slurry for the respective ubiquitin IPs were loaded. The gels were run in 1 x MOPS (Invitrogen) at 180 V for 50 min.

Subsequently, the protein was transferred onto 0.45 µm TransBlot Turbo nitrocellulose (Bio-Rad) membrane using the manufacturer’s High MW setting of the TurboBlot (Bio-Rad). Once the transfer was complete, the membrane was blocked in 5 % milk in TBS-T (0.02% Tween-20) for 1 hr at room temperature while shaking. Next, the membranes were incubated with the primary antibody using a 1:1000 dilution in 5 % TBS-T (monoclonal anti-puromycin antibody, clone 12D10 (Sigma-Aldrich), monoclonal anti-ubiquitin antibody (A-5), sc-166553 (Santa Cruz)). The incubation of the primary antibody was carried out overnight at 4°C while constantly shaking. Afterwards, the membranes were washed three times with TBS-T. The secondary anti-mouse antibody (Millipore, AP181P) with a dilution of 1:10,000 in TBS-T was applied for 1 hr at room temperature while shaking. Then, the membrane was washed three times in TBS-T. The membranes were developed using the Clarity Max Western ECL solution (BioRad), and were analyzed using the Chemidoc (BioRad).

## Computational analysis of ribosome profiling data

### Data pre-processing

A data processing pipeline was developed using Snakemake ^62^ to pre-process raw sequencing reads (.fastq files) and to generate normalized ribosome profiling count tables. This workflow was applied to both monosome and disome datasets, with parameters adjusted accordingly to define the expected A-site position on the leading ribosome.

The workflow proceeded as follows: raw reads were first trimmed to remove Illumina adapter sequences and, if applicable, a 4-nt random sequence. For the DisomeSeq dataset, paired-end reads were merged using “fastp --merged” and subsequently treated as single-end reads. The merged reads were then processed using a published ribosome profiling workflow by Galmozzi et al.,^52^ which includes steps for the removal of reads mapping to noncoding RNAs, A-site mapping, and read counting.

While adapting this workflow, all parameters were kept at their defaults, except for adjustments made to the range of fragment sizes and the positioning of A-sites for each fragment size, similar to Arpat et al.^27^ These adjustments were guided by the observed 3-nt periodicity distributions (**Supplementary Fig. 1d**). Specifically, for monosome data, A-site positions were defined as nucleotide positions {17,18,19}, {18,19,20}, {18,19,20}, {19,20,21} for fragment sizes of 30 to 33 bp. For disome data, the A-site positions for the leading ribosome were defined as {47,48,49}, {47,48,49}, {48,49,50}, {47,48,49}, {50,51,52}, {51,52,53}, {51,52,53} for fragment sizes of 59 to 65 bp.

The resulting raw count tables were subsequently converted into codon-wise count tables by summing up read counts per codon position. To allow comparisons across different experiments, count tables were normalized to a unified library size. Total, Ub-IP, and mock control samples were normalized together, while day 0 and day 3 time points were normalized independently, because of the substantial changes in ribosome occupancy associated with aging. Library size factors were calculated using the “median-of-ratios” method implemented in the DESeq2 package,^63^ based on read counts summed at the gene level. These factors were then applied back to the codon-wise count tables to obtain normalized read counts for downstream analysis.

### Pearson correlation

A Pearson correlation coefficient matrix was computed to include all pairwise correlations between samples with unique identifiers (disome/monosome, total/IP/mock, replicate number). The coefficients were calculated based on signal intensities detected at the codon level, using the normalized codon-wise count table (see the ‘Data Pre-processing’ section above).

### Disome and RQC site identification

Total disome reads were assessed by consistency and expression thresholds. In brief, disome sites were omitted if less than 3 out of 4 experiments contained quantifiable reads, in addition to a mean read cutoff of 10. These sites were then used to perform DESeq2 analysis of Ub-IPs *versus* mock in a codon-wise manner. To estimate log_2_FoldChanges, we applied the ‘ahsr’ log_2_FoldChange shrinkage package^64^ included in DESeq2, to reduce overestimation of significant sites. Significantly enriched sites must exceed a log_2_FoldChange threshold of ≥ 1 and an adjusted *p*-value ≤ 0.05.

Assessment of proximal (59-61 nt) and distal (63-65 nt) disomes was carried out on the disome sites identified in total disome from the whole disome sequencing data set (59-65 nt). We then determined total proximal and distal disome sites fulfilling our consistency and expression cutoff. These sites were then used to run the DESeq2 analysis of the Ub-IP/mock in analogy to the whole disome sequencing data set (*see above*).

### Pause score calculation and pause site identification

Pause scores were calculated in analogy to Stein et al.^16^ Briefly, monosome profiling data were filtered for ≥64 reads per gene. To calculate the pause scores for each gene, we first excluded the first and last 20 codons to exclude start and stop codons. The pause score for each position was computed by dividing the reads for a position within a gene by the average read count for that gene per replicate. The mean pause score for each position was then calculated by averaging across replicates. Pause sites were considered only if the pause score ≥ 2 and contained ≥10 reads. Subsequently, pause sites were subjected to DESeq2 analysis as described for disomes (*see above*).

### Nascent protein identification

The codon-wise read count table was deconvoluted using a sliding window approach, applying a window size of 28 codons with a step size of 14 codons. Read counts within each 28-codon window were summed to represent the aggregated signal for that window. Differential analysis was then performed to identify ubiquitination signals by comparing counts for PAN, K63, and K48 against their respective mock controls within each codon window. This analysis was conducted using the DESeq2 workflow with default parameters, employing false discovery rate (FDR) correction to account for multiple testing.

After DESeq2 analysis, nascent chains were identified using following filters: (*i*) we retained codon windows exceeding a log_2_FoldChange ≥ 1 in CHX and puro and had an adjusted *p*-value ≤0.05 in the CHX experiment; (*ii*) we filtered for codon windows in which log_2_FoldChange(CHX) > log_2_FoldChange(puro); (*iii*) maintained genes with ≥ 6 codon windows and where the mean(log_2_FoldChange(CHX)) > mean(log_2_FoldChange(puro)).

Finally, we applied a sigmoidal fitting on the Ub-IP to mock ratios in a codon-wise manner (*iv*). To analyze the ratio, a 4-parameter logistic (LL.4) model was fitted to the smoothed data using the ‘drc’ package in R. Before fitting, data were smoothed using LOESS regression with a span of 0.4 and a polynomial degree of 2, followed by bootstrap resampling (1000 iterations) to calculate 95% confidence intervals for the smoothed values.

The fit was evaluated based on two criteria: (*i*) the ratio must exceed fold change ≥ 2 at any position along the gene, and (*ii*) the overall trend must show an incline, meaning the final ratio should be greater than the initial ratio. Genes that met these criteria were classified as hits. Genes considered ‘manually curated hits’ are genes with a clearly identifiable onset in their LOESS-smoothed ratio (fold change (Ub-IP *versus* mock) ≥ 2).

### Uniform Manifold Approximation and Projection (UMAP) of UbSeRP multimodal data

UbSeRP data were filtered before dimensionality reduction. Monosome datasets (CHX- and puromycin-treated) were filtered using the DESeq2 calculated baseMeans (threshold≥ 20). All features were subjected to log_2_(fold change) and adjusted *p*-value cutoffs (log_2_(fold change) ≥ 1 and *padj*≤ 0.05), with monosome puromycin features being retained when the matched monosome CHX feature was significant. For pause sites and disomes, only a single feature per window was considered, defined as the site with the maximum log_2_(fold change) within the corresponding monosome window. Windows lacking any significant feature were removed.

UMAP was performed using the uwot package with cosine distance, 80 neighbors, and a minimum distance of 0.4. The resulting two-dimensional embedding was clustered by *k*-means (*k*=3). For visualization, points were colored either by cluster assignment or by the log_2_(fold change) value of the individual datasets.

### Comparison of UbSeRP with MPRA data

MPRA data for RQC were obtained from Chen et al.^29^ To reduce the complexity of the dual RQC reporter sequences, codons were converted to di-amino-acid combinations and then grouped into biochemical class combinations, defined by the properties of the two amino acids (basic, acidic, hydrophobic, polar or stop; for example, basic-hydrophobic). Mean log_2_(fold changes) were calculated for each class combination in wildtype and *hel2*11 samples using the Supplementary Information provided in Chen et al.^29^ Differences between genotypes were assessed using two-sided *t*-tests with Benjamini–Hochberg correction.

To compare these MPRA-derived patterns with endogenous non-ubiquitinated and ubiquitnated disome sequences, 16-amino acid windows upstream of total disomes and PAN- or K63-ubiquitinated disomes were extracted. These sequences were then converted into the same reduced biochemical class representation. Similarity to the MPRA-derived class patterns was quantified using the Hamming distance, and each endogenous sequence was assigned to the closest matching biochemical pattern. Enrichment of these patterns was then calculated for ubiquitinated disomes relative to total disomes. Finally, the change in MPRA signal between *hel2*11 and wildtype was compared with the enrichment of the corresponding biochemical patterns in PAN- and K63-Ub disomes *versus* total disomes.

### Amino acid and codon enrichment analysis

To assess amino acid and codon enrichment within disome and pause sites, we first mapped all total disome and pause sites, as well as significantly enriched PAN and K63-ubiquitinated disome and pause sites, to their corresponding A site codons, amino acids, and genes. These sites were then mapped to their respective open reading frames (ORFs), and the P and E sites were determined.

For each site, the observed frequency of distinct codons and amino acids was calculated. The background distribution was derived from the frequency of each codon and amino acid found in the genes containing either disomes or pause sites. The enrichment value was then calculated as the ratio of observed to expected frequencies.

Statistical significance of the enrichment was assessed using Fisher’s exact test, comparing amino acid or codon occurrences within the observed distribution to those in the background distribution. Multiple testing correction was performed using the Benjamini-Hochberg method. Unless otherwise stated, only sites with a log_2_FoldChange ≥ 0.5 and adjusted *p*-value ≤ 0.05 were considered in the analysis to identify significantly enriched amino acids at the A, P, and E sites.

### Metagene analysis of biophysical parameters

To investigate local biophysical properties around disome and pause sites, we used distinct filtering strategies to exclude ubiquitination-dependent events. For disomes, we first clustered all high-confidence sites (total disomes) by gene and genomic position, grouping adjacent sites into clusters. Clusters containing any significantly enriched PAN-Ub disomes were excluded in their entirety to avoid contamination from ubiquitin-associated signals. In contrast, for pause sites, we directly removed any positions overlapping significant PAN-enriched events without clustering. For each remaining gene, 100 random positions were sampled to serve as background. Amino acid windows spanning –40 to +10 codons relative to the A-site were extracted for both observed and random positions. Biophysical parameters, including hydropathicity,^65^ isoelectric point (pI; https://www.sigmaaldrich.com/DE/de/technical-documents/technical-article/protein-biology/protein-structural-analysis/amino-acid-reference-chart), amino acid volume,^66^ and codon optimality^67^ were computed at each relative position. For codon optimality, we used a binary system: optimal = 1 and non-optimal = 0. Background values were averaged per gene and position, and observed windows were normalized by subtracting these gene-specific means. Metagene plots display the mean difference between observed, and background signals, and 95% confidence intervals were calculated across genes. To assess the significance of total *versus* PAN-Ub sites, the area under the curve (AUC) from –20 to 0 codons was computed and tested using a Wilcoxon rank-sum test.

### Correlation of 5’-3’ co-translational degradation with disomes (related to Fig. 3j)

To correlate 5’-3’ co-translational degradation with disomes, we obtained previously published codon protection indices from Pelechano et al.^34^ Codon protection indices were stratified into four bins (‘low’ (most stable), ‘medium,’ ‘high,’ ‘very high’ (most unstable)) based on quartiles. Disome frequency was calculated per gene as the number of mapped disome footprints normalized to the median disome count across all genes. Ubiquitination frequency was calculated as the number of PAN-ubiquitination events per gene divided by the total number of disome footprints for that gene. Genes with a ubiquitination frequency of 0 were excluded from the comparison. Disome and ubiquitination frequencies were compared across the codon protection index bins using pairwise Wilcoxon rank-sum tests.

### Enrichment of structural features at disome sites

Protein domain, coiled-coils, and zinc fingers information was obtained from UniProt (https://www.uniprot.org/), transmembrane domains (TMDs) were predicted using TMHMM v2.0 (https://services.healthtech.dtu.dk/services/TMHMM-2.0/), and chaperone binding sites for Ssb1, Ssb2, and TRiC for genes with disome sites were identified based on published data from Döring et al. and Stein et al.^40,41^

Disome enrichment was quantified across structural and functional features by comparing total disomes to PAN-Ub disomes. For annotated protein domains, the exact domain boundaries from UniProt were used to define the overlap of domains with total and PAN-Ub disomes. For transmembrane domains (TMDs), a +30 to +80 codon window downstream of the annotated TMD start site was defined to capture co-translational stalling near membrane insertion signals. Chaperone association was assessed using binding site data for Ssb1, Ssb2, and TRiC, by defining a –20 to +10 codon window around each disome site and detecting overlaps with chaperone-binding peaks. For each feature, the odds ratios were calculated from contingency tables comparing the frequency of PAN-Ub versus total disomes within versus outside the feature window, and statistical significance was assessed using Fisher’s exact test.

### Gene ontology analysis

GO-term analysis for the biological process (bp), molecular function (mf), and cellular component was performed using STRINGdb v12.0.^68^ The resulting GO-terms were manually curated with an emphasis on gene expression, metabolism, and protein folding to reduce redundancy of terms. A full list of GO-terms will be made available upon publication.

### Analysis of co-occurrence of ubiquitination with chaperone binding, degrons and structural features

Nascent protein ubiquitination sites were analyzed for overlap with chaperone binding regions, predicted degron features, and annotated structural domain. Chaperone binding sites for Ssb1, Ssb2, and TRiC were extracted from supplementary datasets provided in ^40,41^. Degron predictions and motifs were obtained using Degronopedia v2.1.0,^39^ and structural features were retrieved from UniProt (https://www.uniprot.org/).

Overlaps were determined based on the alignment of ubiquitin-modified regions with annotated feature intervals. Chaperone associations for Ssb1 and Ssb2 were analyzed within ±40 codons of ubiquitination sites, and overlap proportions were visualized. Degron motifs were located in a −80 to −30 codon window upstream of the nascent chain ubiquitination onset were considered.

### Random Forest Classifier

To identify sequence features predictive of RQC targeting, we trained a Random Forest binary classifier to distinguish RQC sites (stalled ribosomes detected in both Disome-PAN and Disome-Total fractions) from non-RQC sites (detected in Disome-Total only). Features comprised physicochemical properties of a 50-amino-acid window (isoelectric point, bulkiness, hydrophobicity, alpha-helix and beta-sheet propensity, and flexibility), structural context (number of overlapping transmembrane domains and annotated protein domains), co-translational chaperone sites (Ssb1 and TRiC), individual codon frequencies, codon optimality (fraction of optimal codons), and amino acid frequencies. Model performance was evaluated by repeated stratified 10-fold cross-validation (10 repeats, 100 folds total; “RepeatedStratifiedKFold” from scikit-learn package), with class imbalance handled by inverse-frequency sample weighting (class_weight=’balanced’). The model was trained as follows: physicochemical properties within the sequence windows were corrected by the background of same-size windows along the corresponding protein. Recursive Feature Elimination (RFE, step = 1) was applied across 10 stratified folds to rank features by their contribution to classification performance.

### Comparison of differential translation in aging

To assess gene-specific translational dynamics during aging, normalized expression scores were derived from total monosome profiling across multiple time points (day 0, day 2, and day 3). Genes with an average read count ≤ 32 across all conditions were removed. For each remaining gene, the mean and standard deviation of expression across all days were computed, and to obtain a z-score in a gene-wise manner across aging:

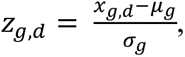

where *z_g,d_* is the normalized score for the gene *g* on day *d*, *x_g,d_* is the expression of gene *g* on day *d*, *µ_g_* is the mean expression of the gene across all days and *α_g_* is the standard deviation. These z-scores were visualized in a heatmap. Rows were clustered using hierarchical clustering based on Euclidean distance, and genes were divided into three clusters to capture distinct temporal expression profiles. Subsequently, GO-term analysis was carried out in a cluster-wise manner (see **“Gene ontology analysis”**).

### Assessing the contribution of codon optimality to age-dependent amino acid bias in ribosomal sites

To test whether codon optimality explains differences in enrichment ratios between day 3 and day 0 at the E, P, and A ribosomal sites, we modeled each codon as an independent observation with recorded enrichment scores at both time points. A baseline linear regression model was first fitted as the following formula (in R format):

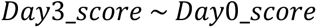

To capture the correlation between the two time points. We then extended this to a “complex” model including codon optimality as a binary predictor:

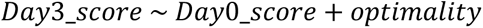

Codon optimality was encoded as a binary variable. The additional contribution of codon optimality was assessed using an ANOVA F-test, using the ‘anova()’ function in R, comparing the baseline and complex models (equivalent to a t-test in this single-parameter case). Statistical significance indicated that codon optimality explained additional variation in day 3 enrichment scores beyond day 0 scores.

### AlphaFold3 structural predictions

Structures shown in **Supplementary Fig. 6c** were generated using AlphaFold3.^69^ Structures were analyzed using ChimeraX.^70^

### Data visualization

Average read counts were calculated for each condition (e.g., PAN, K48, K63) and biological replicate, considering only positions with at least three non-zero values across replicates. Ratios of treatment to total reads were computed for each position, representing the treatment condition divided by the corresponding total condition (e.g., CHX-PAN/total, puro-PAN/total).

LOESS smoothing was applied to the ratio data with a span of 0.2 and a polynomial degree of 2. Bootstrap resampling (1000 iterations) was used to calculate 95% confidence intervals, based on the 2.5th and 97.5th percentiles. The smoothed data and confidence intervals were visualized as a solid line and shaded area, respectively.

### Quantification and statistical analysis

Significance was denoted as follows: n.s., *p* > 0.05; \**p* ≤ 0.05; \*\**p* ≤ 0.01; \*\*\**p* ≤ 0.005; \*\*\*\**p* ≤ 0.001. *p*-values were adjusted for multiple testing using the Benjamini-Hochberg procedure. All statistical tests were two-sided unless otherwise specified. For disome-seq heatmaps, asterisks indicate codons that are statistically significant, independent of the significance level annotation used elsewhere.

UbSeRP experiments were performed with *n*= 4 biologically independent replicates. Unless stated otherwise, an enrichment threshold of log_2_(fold change)≥ 2 was applied (for aging experiments, log_2_(fold change)≥ 1.5); specific thresholds are indicated in the corresponding figure legends.

Western blot analyses were conducted with at least *n*= 2 biologically independent replicates.

## Supplementary Figures

**Supplementary Figure 1.**
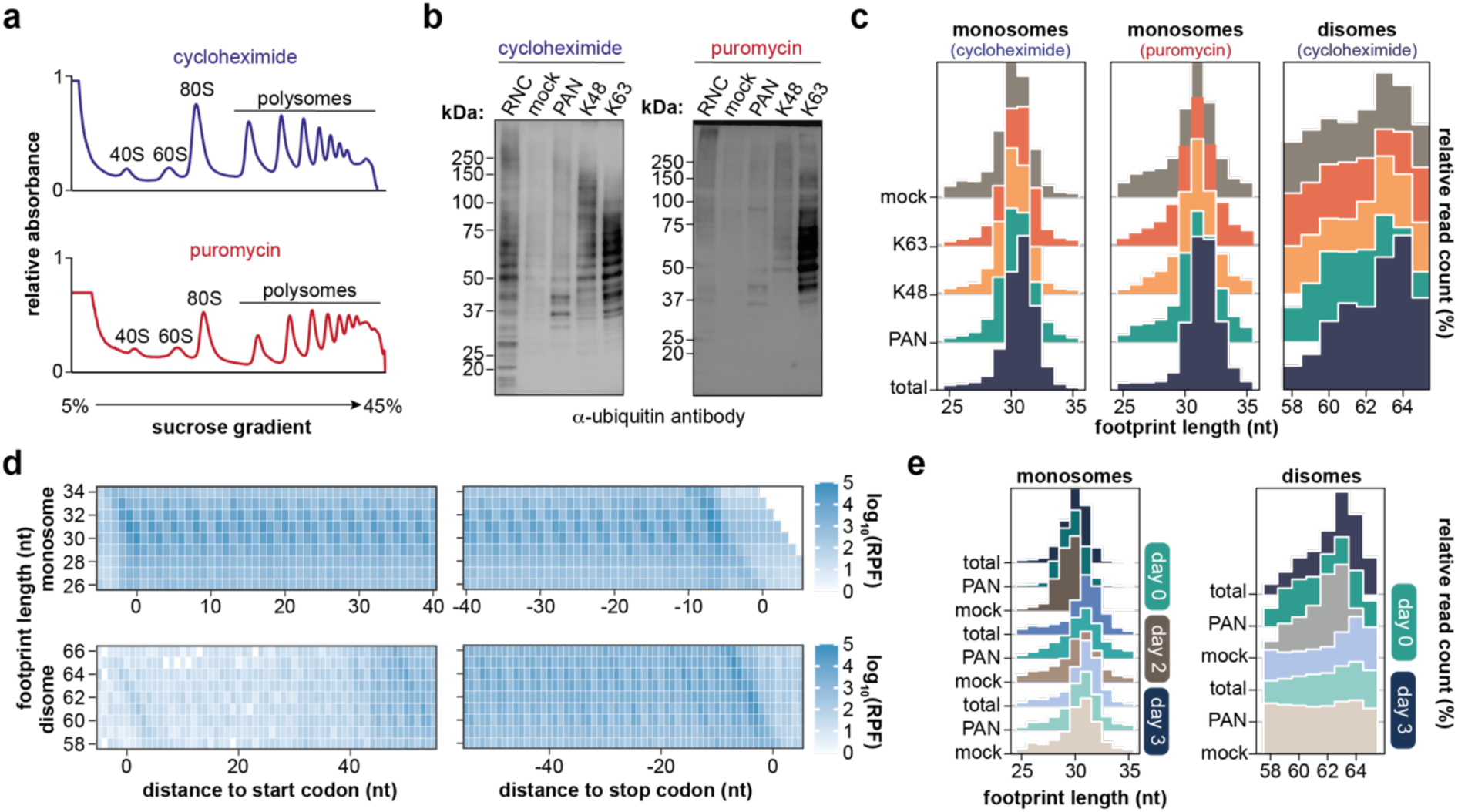
Quality control of UbSeRP experiments. **a,** Representative polysome profiles of cycloheximide-treated (*top*) and puromycin-treated (*bottom*) lysates indicate the integrity of polysomes under our lysis conditions. **b,** Representative Western Blot analysis of ribosome-nascent chain complexes (RNCs; input) and the corresponding TUBE-enriched ubiquitin eluates from cycloheximide- and puromycin-treated RNC samples and their respective immunoprecipitations showing the accumulation of ubiquitinated proteins in TUBE pulldowns as compared to mock controls. Mock samples correspond to lysates incubated with streptavidin beads to estimate background arising from biotinylated contaminants. Ubiquitin was detected using an anti-ubiquitin mouse antibody. **c,** Ribosome footprint length distributions for monosome and disome profiling experiments used to investigate ubiquitin linkages. Distributions show relative read counts across four biologically independent experiments (*n=*4). **d,** Heatmap showing the 3-nt periodicity for total monosome (*top*) and disome (*bottom*) profiling datasets. Plots show strong 3-nt periodicity at the start and stop regions of open reading frames (ORFs). Color scale indicates reads per ribosome-protected fragment (RPF). **e,** Same as in **c**, but for monosome (*left*) and disome (*right*) profiling datasets from the aging experiments.

**Supplementary Figure 2.**
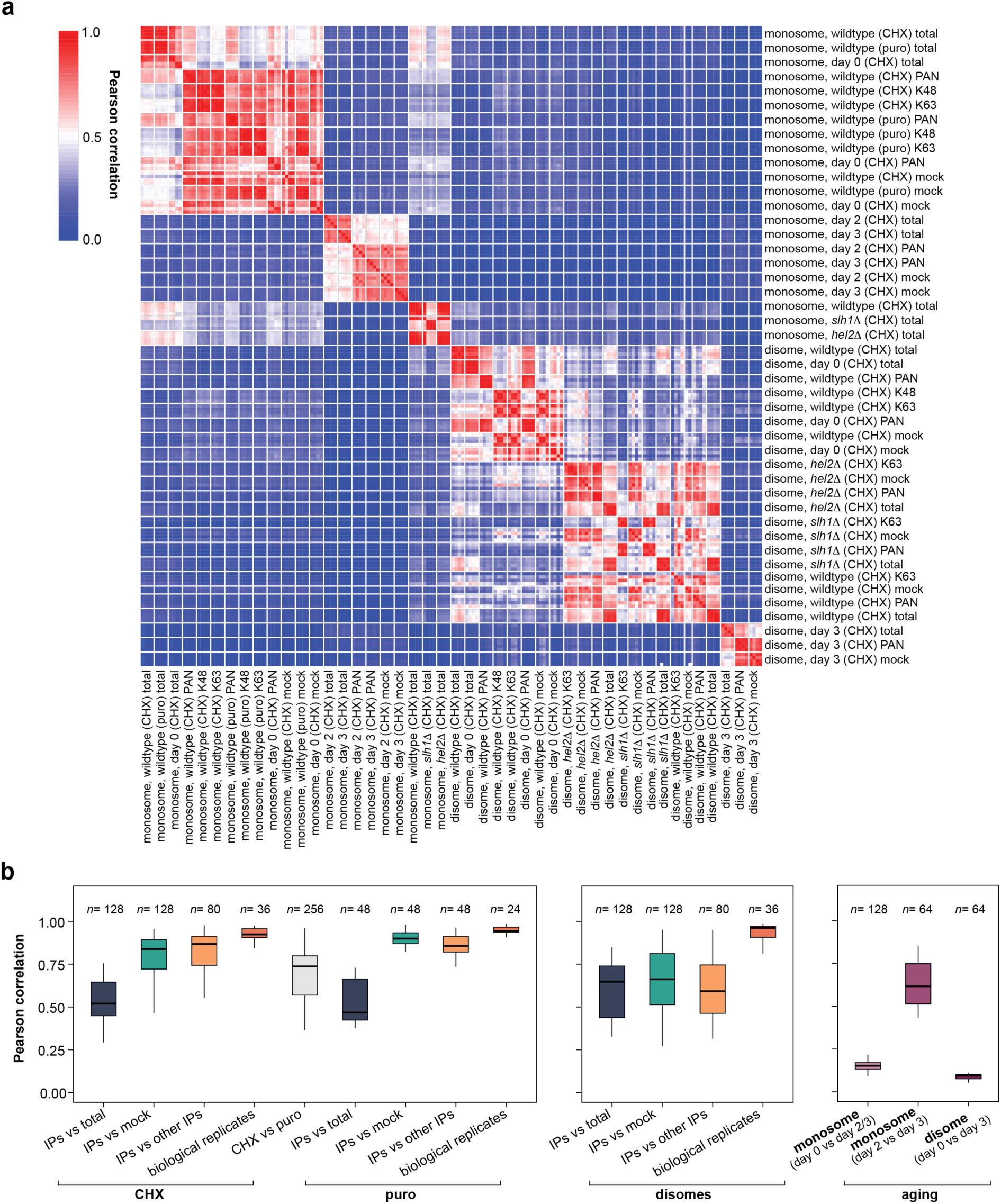
Assessment of data consistency. **a,** Heatmap of the Pearson correlation of the selective mono- and disome experiments for all experiments presented in this manuscript. Matrix displays the correlation based on normalized read count at single-codon resolution. **b,** Box plots summarizing Pearson correlation across all experiments. Comparisons are grouped, for example, IPs (PAN, K48, and K63) *versus* total across all four biological replicates (*n*=4) in wildtype backgrounds. Self-correlations are omitted. Pearson correlation coefficients are based on **a**. The box plots present the median as the central solid line, the bottom and top edges of the box as the IQR, and the box plot whiskers represent 1.5 times the IQR.

**Supplementary Figure 3.**
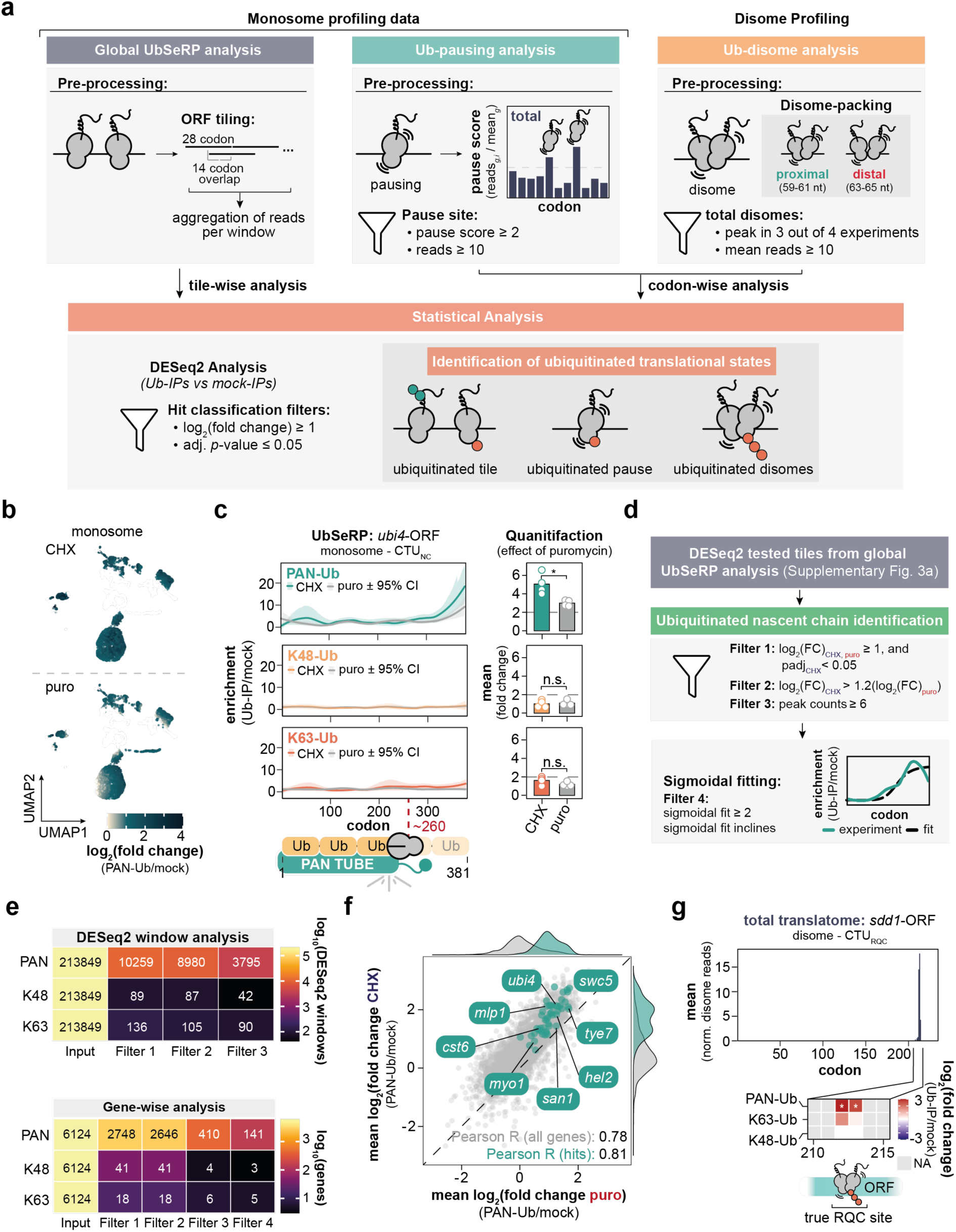
Ubiquitination features during translation. **a,** Schematic overview of the data-processing pipelines used to derive ubiquitination-associated features during translation; details are provided in the **Methods**. Monosome and disome profiling datasets were processed separately to capture global ubiquitination patterns, translational pausing, and disome formation (*top*). The resulting outputs were then used for DESeq2 analysis of Ub-IP enrichment by comparing ubiquitin immunoprecipitations (Ub-IPs) with mock controls (*bottom*). **b,** UMAP representation of UbSeRP data. Data points are colored according to log_2_(fold change) values calculated for PAN-Ub versus mock in cycloheximide (CHX)- and puromycin (puro)-treated samples (DESeq2 results from the global UbSeRP analysis). The color scale was capped between 0 and 4. **c,** Representative selective ribosome profiles for *ubi4*-ORF from PAN-Ub (*top left*), K48-Ub TUBE (*middle left*), and for K63-Ub TUBE pulldowns (*bottom left*) with corresponding quantification (*right*). Nascent Ubi4 is enriched at codon ∼260 in the PAN-Ub pulldown, but not in the K48- and K63-Ub pulldowns. Profiles show LOESS-smoothed enrichment (PAN-, K48- or K63-Ub versus mock non-binding control) from CHX- and puro-treated ribosome profiling experiments across four biologically independent replicates (*n=*4). Shaded areas indicate bootstrapped 95% confidence intervals. Statistical significance was assessed using a two-sided paired *t*-test comparing cycloheximide- and puromycin-treated samples: PAN-Ub versus mock, \**p* = 0.043; K48-Ub versus mock, *p* = 0.79; K63-Ub versus mock, *p* = 0.19. **d,** Workflow of the detection pipeline. DESeq2 results of the global UbSeRP analysis were taken as input and subsequently filtered by log_2_(fold changes) in both, CHX and puromycin, differential expression in CHX and puromycin, and the number of peak counts within each respective gene. Short-listed genes were subjected to sigmoidal fitting using the unbinned ribosome profiling data. **e,** Heatmap of the effect of filters (review panel **d**) on UbSeRP data for analysis windows (28 codons with 14 codons sliding window; *top*) and per gene (*bottom*). **f,** Pearson correlation of mean log_2_(fold change) per gene for CHX and puro. **g,** Bar plot showing mean normalized total disome reads for *sdd1*-ORF. The heatmap below shows Ub enrichment based on DESeq2 analysis. The true-positive RQC site corresponds to the previously described RQC site in *sdd1*-ORF. The log_2_(fold change) scale ranges from -3 to 3, and significant codons are indicated by asterisks (*padj* ≤ 0.05). NA indicates codon positions for which log_2_(fold change) could not be determined due to low reproducibility or expression cutoffs. Data are derived from four biologically independent UbSeRP experiments per condition (*n=*4). CHX: cycloheximide, ORF: open reading frame; puro: puromycin; TUBE: tandem ubiquitin binding entity; Ub: ubiquitin; FC: fold change. n.s., *p* > 0.05; \**p* ≤ 0.05; \*\**p* ≤ 0.01; \*\*\**p* ≤ 0.005; \*\*\*\**p* ≤ 0.001.

**Supplementary Figure 4.**
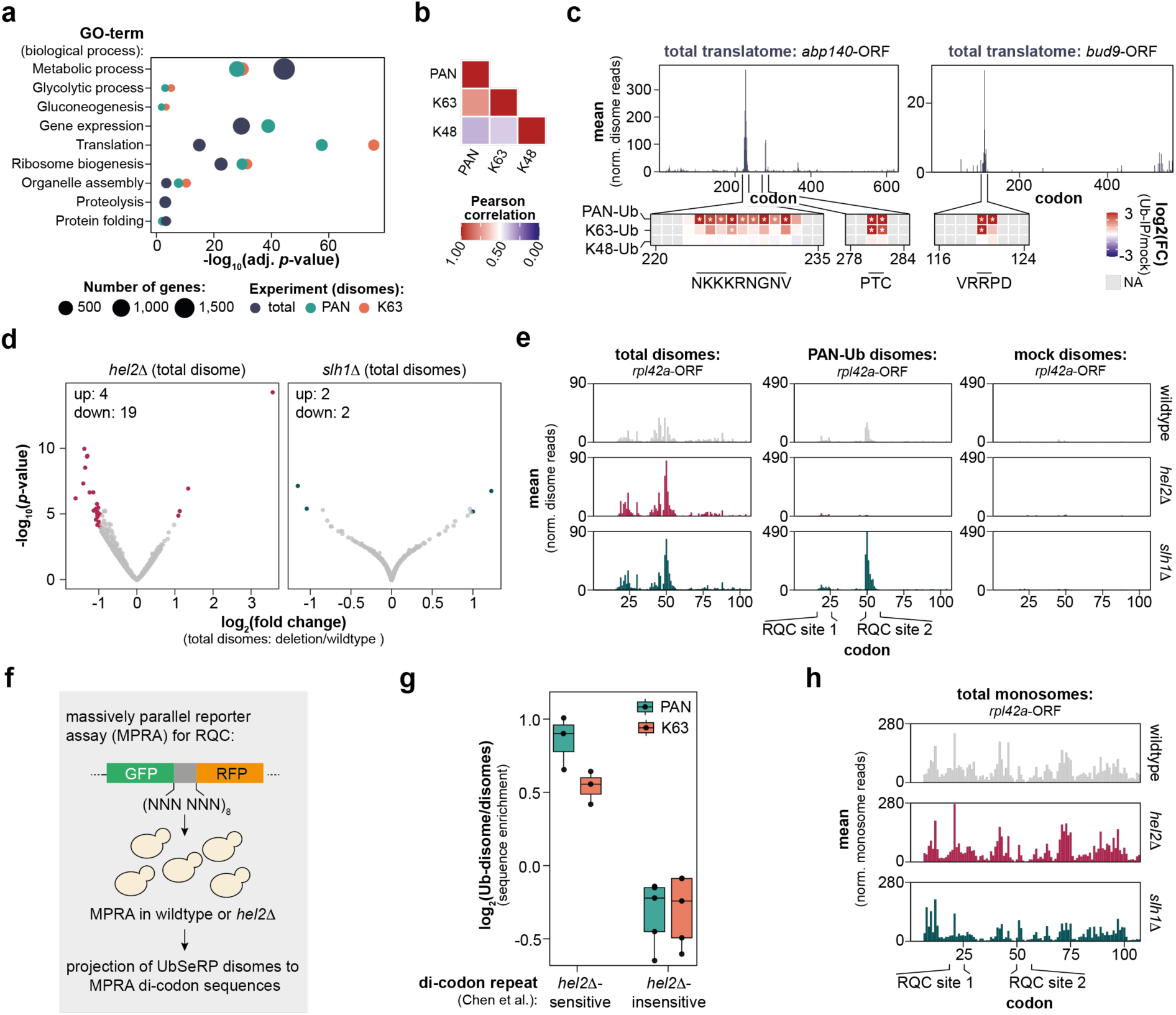
Characterization of UbSeRP disome profiling. **a,** Gene ontology (GO) analysis of biological process terms for genes containing disomes. GO terms were manually curated, with a focus on metabolism, gene expression, and proteostasis. **b,** Pearson correlation of log_2_(fold change) values for PAN-Ub hits across all pairwise comparisons between PAN-, K63-, and K48-Ub enrichments. **c,** Representative total disome profiles for *abp140*-ORF and *bud9*-ORF. Bar plot showing mean normalized total disome reads. The heatmap below shows Ub enrichment based on DESeq2 analysis. The log_2_(fold change) scale ranges from -3 to 3, and significant codons are indicated by asterisks (*padj* ≤ 0.05). NA indicates codon positions for which log_2_(fold change) could not be determined owing to expression cutoffs. Data are shown from four biologically independent experiments per condition (*n*=4). **d,** Volcano plot for total disomes comparing RQC knockouts *versus* wildtype. Hits exceeding the log_2_(fold change) and adjusted *p*-value cutoff are indicated (log_2_(fold change) ≥ ±1; *padj* ≤ 0.05) as colored dots and non-significant codons as grey dots. Statistics were computed using DESeq2. **e,** Representative disome profiles for *rpl42a*-ORF showing total, PAN-Ub, and mock disomes in wild type, *hel2*Δ, and *slh1*Δ cells. Bar plot showing mean normalized total disome reads. Data are shown from four biologically independent experiments per condition (*n*=4). **f,** Schematic of the massively parallel reporter assay (MPRA) published by Chen *et al.*^29^ and the associated data-analysis workflow. **g,** Sequence enrichment analysis of PAN-Ub and K63-Ub disomes categorized as *hel2*Δ-sensitive or *hel2*Δ-insensitive based on the MPRA dataset. **h,** same as in **e**, but for monosome profiling data for *rpl42a*-ORF.

**Supplementary Figure 5.**
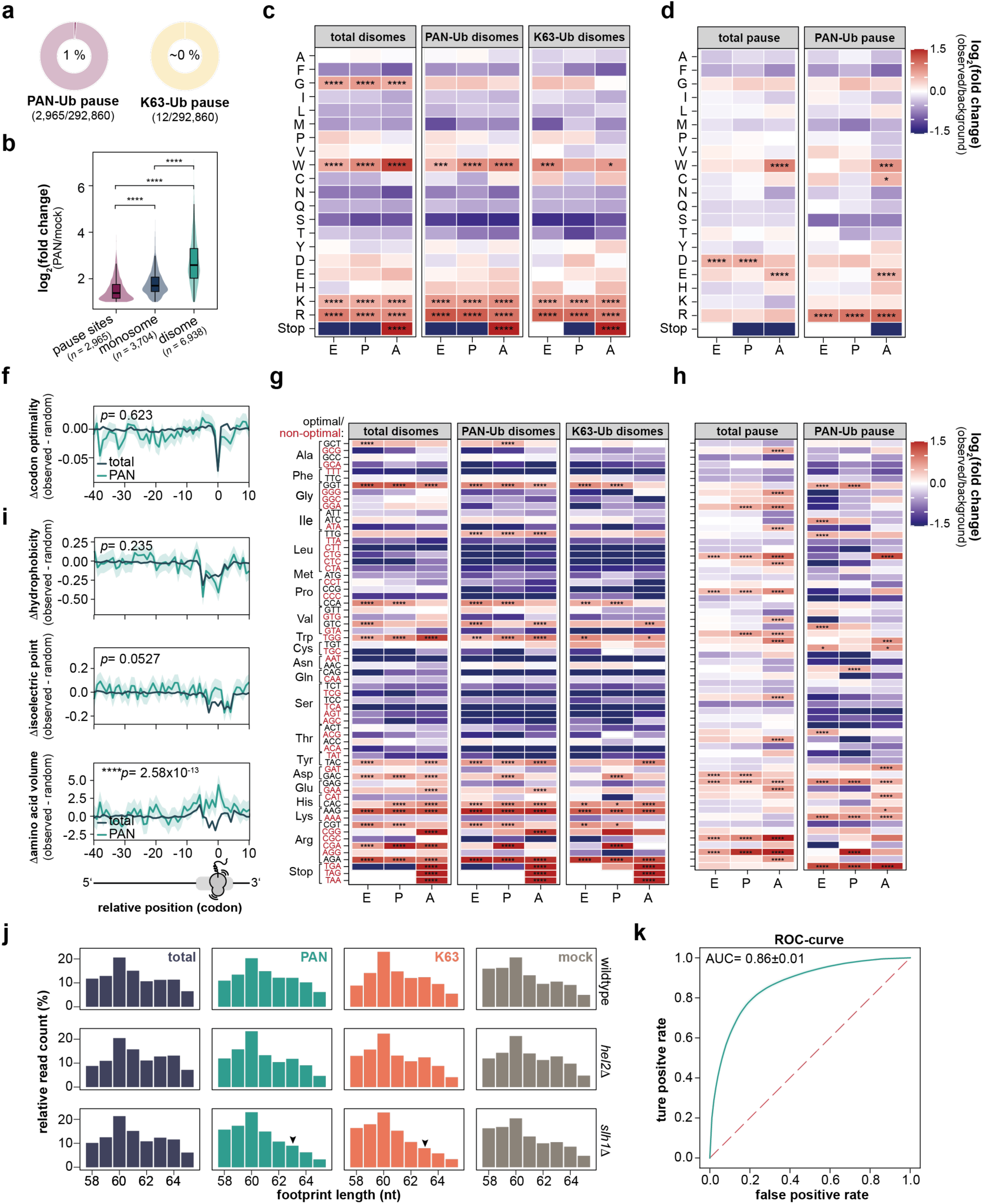
Mechanistic insight into CTU_RQC_. **a,** Pie chart showing the polyubiquitination frequencies of PAN-Ub and K63-Ub pause sites. **b,** Comparison of PAN-polyubiquitinated pause sites, monosome-projected disomes, and disomes. Statistical significance was assessed using the Wilcoxon test: \*\*\*\**p* = 1.93 × 10^-149^ (pause sites *versus* monosomes), \*\*\*\**p* = 0 (pause sites *versus* disomes), and \*\*\*\**p* = 0 (monosomes *versus* disomes). **c,** Heatmap showing amino acid bias at the exit (E), peptidyl transferase (P), and aminoacyl (A) sites for total, PAN-Ub, and K63-Ub disomes. The color scale indicates log_2_(fold change) of observed versus background bias. Adjusted *p*-values are indicated by asterisks (Fisher’s exact test with Benjamini-Hochberg correction). **d,** Same as in **c**, but for total and PAN-Ub pause sites. **e,** Metagene plot of codon optimality at total and PAN-Ub disome sites. The solid line indicates the mean Δcodon optimality at disome sites, and the shaded area represents the 95% confidence interval. No significant difference was observed (Wilcoxon test, n.s., *p*= 0.623). **f,** and **g,** heatmap of codon-level analysis of disome (**f**) and pause-site (**g**) datasets for total and PAN-Ub signals. Non-optimal codons are shown in red and optimal codons in black. *p*-values were calculated as in **c**. **h,** Metagene plots showing changes in Δhydrophobicity (*top*), Δisoelectric point (pI; *middle*), and Δamino acid bulkiness (*bottom*) at total and PAN-Ub pause sites. In each panel, the solid line indicates the mean change at disome sites and the shaded area denotes the 95% confidence interval. Statistical significance was assessed using the Wilcoxon test: hydrophobicity, *p*= 0.235 (n.s., AUC, positions -10 to 0); pI, *p*= 0.0527 (n.s., AUC, positions -10 to 0); bulkiness, \*\*\*\**p*= 2.58×10^-13^ (AUC, positions -10 to 0). **j,** Disome footprint length distributions for disome profiling experiments for wildtype, *hel2*Δ and *slh1*Δ. Distributions show relative read counts across four biologically independent experiments (*n*=4). The arrow head indicates the depletion of distal disomes in *slh1*Δ background in PAN- and K63-Ub IPs. **k,** Receiver operating characteristic (ROC) curve showing recovery of RQC events among all identified disomes. The shaded area indicates ±1 standard deviation across 100 cross-validation folds (10 repeats for 10-fold stratified CV). The red dashed line indicates the random guessing. n.s., *p* > 0.05; *\*p* ≤ 0.05; *\*\*p* ≤ 0.01; *\*\*\*p* ≤ 0.005; *\*\*\*\*p* ≤ 0.001.

**Supplementary Figure 6.**
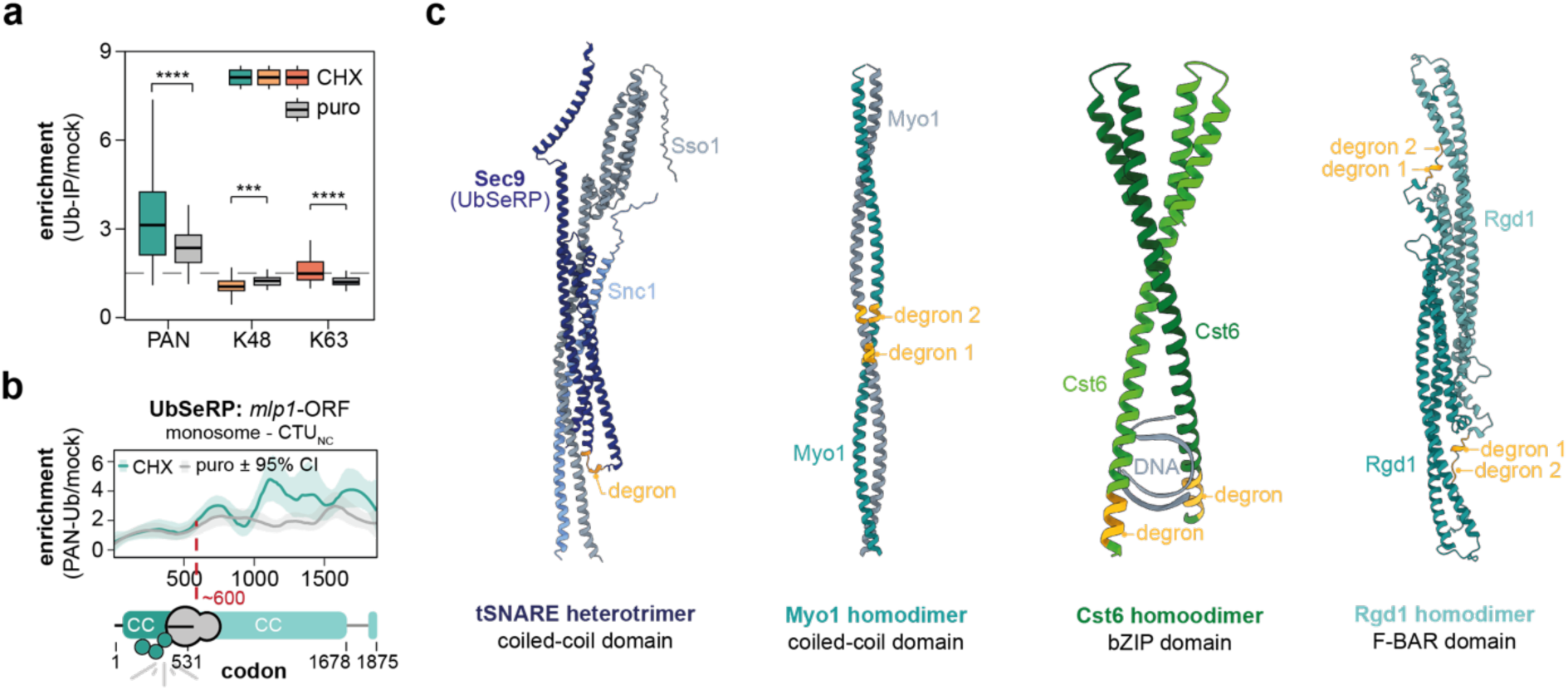
Development and benchmarking of a nascent chain detection framework for UbSeRP. **a,** Box plot for the assessment of puromycin-sensitivity and effect size of PAN-, K48-, and K63-linked polyubiquitinated nascent chains. The box plots present the median as the central solid line, the bottom and top edges of the box as the IQR, and the box plot whiskers represent 1.5 times the IQR. Pairwise comparisons of the enrichment for cycloheximide (CHX) or puromycin (puro) treated Ub-enrichments (Ub-IP *versus* mock) were performed using the Wilcoxon test: PAN-Ub, \*\*\*\**p* = 8.46 × 10^-8^; K48-Ub, \*\*\**p* = 1.23 × 10^-3^; K63-Ub, \*\*\*\**p* = 3.01 × 10^-7^. The grey dashed line indicates 1.5-fold enrichment for *n*= 144. **b,** Representative selective ribosome profiles for *mlp1*-ORF from a PAN-Ub TUBE pulldown. Profile shows LOESS-smoothed enrichment (PAN-Ub versus mock non-binding control) from CHX- and puro-treated ribosome profiling experiments across four biologically independent replicates (*n*=4). Shaded areas indicate bootstrapped 95% confidence intervals. **c,** Representative examples of proteins targeted by CTU_NC_ in their native complexes. Predicted degrons corresponding to the onset are highlighted in orange. Structures were generated using Alphafold3.^69^ n.s., *p* > 0.05; *\*p* ≤ 0.05; *\*\*p* ≤ 0.01; *\*\*\*p* ≤ 0.005; *\*\*\*\*p* ≤ 0.001.

**Supplementary Figure 7.**
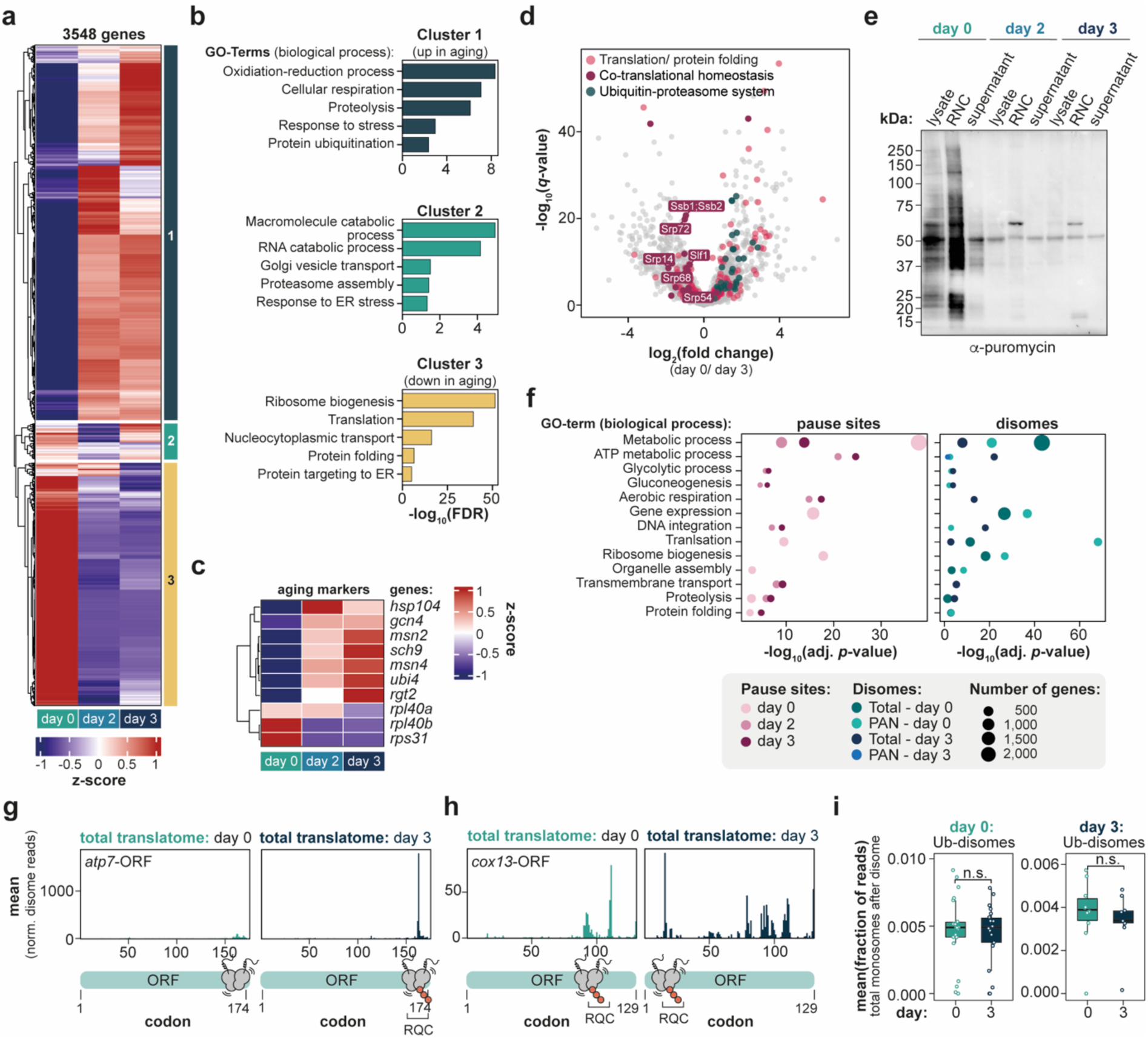
Effects of chronological aging on gene expression and translation. **a,** Heatmap of translated genes, derived from monosome profiling, showing differential expression across aging. Expression changes are displayed as row-wise z-scores for the three time points: day 0, day 2, and day 3. Genes were assigned to one of the three indicated clusters. The heatmap includes 3,548 genes detected with more than 32 reads. The color scale represents z-scores. **b,** Bar plots showing gene ontology (GO) analysis of biological process terms for the three clusters shown in **a**. **c,** Heatmap of aging marker expression across the three time points, displayed as z-scores. *rpl40a, rpl40b*, and *rps31* are ubiquitin genes that are downregulated during cellular stress, whereas *ubi4* is upregulated. **d,** Volcano plot of mass spectrometry-based quantification of ribosome-nascent chain complexes (RNCs), comparing day 0 and day 3. Data are derived from four biologically independent datasets (*n=*4). Annotation of translation/protein folding factors and ubiquitin-proteasome system components is based on the Proteostasis Network Annotation,^71,72^ which we extended to yeast homologs and manual curation to highlight co-translational regulators. **e,** Representative Western Blot analysis of aged cells showing puromycin incorporation in lysates, RNCs, and sucrose cushion supernatants as a measure of translation activity. The experiment was performed in two biologically independent replicates (*n=*2). Puromycin incorporation was detected using an anti-puromycin antibody. Proteins were detected using horseradish peroxidase (HRP)-conjugated anti-mouse secondary antibodies. **f,** GO analysis of biological process terms for genes containing pause sites (*left*), as well as total and PAN-Ub disomes (*right*), across the different aging time points. GO terms were manually curated. **g,** and **h,** Representative total disome profiles for *cox13*-ORF (**g**) and *atp7*-ORF (**h**). Plots show mean total disome reads for day 0 and day 3. Disome icons indicate PAN-Ub and non-ubiquitinated disome sites based on log_2_(fold change) and adjusted *p*-values (*padj* ≤ 0.05). **i,** Box plot quantifying translation, based on monosome profiling, across the 100-codon region downstream of a disome site. Dots represent individual genes and indicate the mean fraction of reads. Pairwise comparisons of the mean fraction of reads were performed using the Wilcoxon test: *left,* day 0 Ub-disomes *versus* day 3 non-Ub disomes, ns, *p*= 0.781 (*n*=22); right, day 3 Ub-disomes *versus* day 0 non-Ub disomes, ns, *p*= 0.154 (*n*=10); see also Fig. 5i. n.s., *p* > 0.05; \**p* ≤ 0.05; \*\**p* ≤ 0.01; \*\*\**p* ≤ 0.005; \*\*\*\**p* ≤ 0.001.

